# Time series transcriptome analysis uncovers regulatory networks and a role for the circadian clock in the *Drosophila melanogaster* female’s response to Sex Peptide

**DOI:** 10.1101/2022.05.14.491957

**Authors:** Sofie Y.N. Delbare, Sara Venkatraman, Kate Scuderi, Martin T. Wells, Mariana F. Wolfner, Sumanta Basu, Andrew G. Clark

**Affiliations:** Department of Molecular Biology & Genetics, Cornell University, Ithaca NY, USA; Department of Statistics & Data Science, Cornell University, Ithaca NY, USA

**Author notes:** Corresponding author: Sofie Y.N. Delbare.

**Keywords:** time-course RNA-seq, gene regulatory network analysis, *Drosophila*, Sex Peptide, circadian rhythm

## Abstract

Sex Peptide, a seminal fluid protein of *D. melanogaster* males, has been described as driving a virgin-to-mated switch in females, through eliciting an array of responses, including increased egg laying, activity and food intake and a decreased re-mating rate. While it is known that Sex Peptide achieves this, at least in part, by altering neuronal signaling in females, the identity of key molecular regulators that act downstream of Sex Peptide is not known. Here, we used a high-resolution time series RNA-sequencing dataset of female heads at 10 time points within the first 24 hours after mating to investigate the genetic architecture, at the gene- and exon-level, of the female’s response to Sex Peptide. We find that Sex Peptide is not essential to trigger a virgin-to-mated transcriptional switch, which involves changes in a metabolic gene regulatory network. However, Sex Peptide is needed to maintain and diversify metabolic changes and to trigger changes in a neuronal gene regulatory network. We further find that Sex Peptide might interact with the female’s circadian clock to orchestrate transcriptional changes across different regulatory networks. That a male seminal fluid protein can alter a female’s rhythmic gene expression has implications for our understanding of both reproductive and circadian behaviors.

## Introduction

While sperm are essential and sufficient for fertilization, males of many animal species, ranging from fruit flies to humans, transfer more than only sperm during mating. Proteins, lipids, vesicles and cells, among others, accompany sperm in the seminal fluid (e.g. Findlay et al. 2008, Pilch and Mann 2006, Leiblich et al. 2012, Corrigan et al. 2014, Conine et al. 2018). Non-sperm seminal fluid components, while not strictly required for fertilization, can increase reproductive success through alterations of the female’s physiology or behavior (Bromfield 2014, Perry et al. 2013, Poiani 2006).In *Drosophila melanogaster*, the seminal fluid protein Sex Peptide (SP) has been shown to be directly causal for a variety of long-term post-mating responses in females (Kubli and Bopp 2012). These responses extend long-term due to SP’s ability to bind to sperm, followed by its gradual release via active proteolytic cleavage. This allows SP to be stably maintained in the female and exert its effects for multiple days (Peng et al. 2005). SP-mediated responses include increased egg laying and reduced receptivity to additional matings (Chapman et al. 2003; Liu and Kubli 2003), and SP is needed to release sperm from storage organs (Avila et al. 2015; Avila et al. 2010). Further, SP leads to an increase in juvenile hormone titers (Bontonou et al. 2015; Moshitzky et al. 1996), immunity changes (Peng et al. 2005; Domanitskaya et al. 2007), alterations to the female’s metabolism, gut morphology and excretion (White et al. 2021; Apger-McGlaughon and Wolfner 2013; Cognigni et al. 2011), decreased sleep during the day (Isaac et al. 2010), an increased appetite, with a preference for high-protein food (Walker et al. 2015; Carvalho et al. 2006; Uchizono et al. 2017; Ribeiro and Dickson 2010), increased aggression (Bath et al. 2017), and an enhanced long-term memory (Scheunemann et al. 2019). A fascinating question is how one protein can induce such diverse effects. Multiple studies have investigated SP’s mode of action to better understand how it controls these post-mating responses.

Most SP-mediated responses require binding of SP to the Sex Peptide Receptor (SPR), a G protein-coupled receptor that is expressed in the female’s reproductive tract and central nervous system (Yapici et al. 2008). Through SPR, SP silences neuronal activity (Yapici et al. 2008). Several studies have started mapping the neuronal circuits that are required to sense SP and to relay SP/SPR signaling. These studies have shown that a common set of neurons, sensory *ppk^+^/fru^+^* neurons that innervate the reproductive tract (Häsemeyer et al. 2009; Yang et al. 2009; Lee et al. 2016), which connect to neurons in the abdominal ganglion (SAG neurons), which in turn project to the pC1 region in the brain (Wang et al. 2020; Jang et al. 2017; Rezával et al. 2014; Feng et al. 2014; Rezával et al. 2012; Soller et al. 2006; Zhou et al. 2014), are all required for SP’s effects on female receptivity (Wang et al. 2021), oviposition (Wang et al. 2020), sleep (Garbe et al. 2016), and feeding behavior (Walker et al. 2015). Together, these studies show that at least some of the post-mating responses induced by SP share a common neuronal basis, and they show that although SP is stored in the female reproductive tract, its signals result in a profound impact on the female’s brain.

In addition to changes in neuronal signaling, SP causes changes in RNA abundance throughout the female body, influencing genes involved in metabolism, development, signal transduction, regulation of transcription, transport, the immune response and phototransduction, based on measurements done in the head/thorax, abdomen and midgut (Gioti et al. 2012; Domanitskaya et al. 2007; White et al. 2021). The gene expression changes identified in these studies likely contribute to phenotypic post-mating responses, but when and how SP taps into gene regulatory networks to mediate these expression changes remains elusive. SP’s fine-scale temporal effects on gene expression have not been studied before, even though we know that gene expression in mated females changes over time (Dalton et al. 2010; Gioti et al. 2012; Mack et al. 2006; Prokupek et al. 2009; McDonough-Goldstein et al. 2021; McGraw et al. 2008), and distinct aspects of mating, such as seminal fluid proteins, sperm or pheromones, exert their effects within different time windows after mating (Heifetz et al. 2014; Rubinstein and Wolfner 2013; McGraw et al. 2004; Tram and Wolfner 1998; Laturney and Billeter 2016).

SP is often described as a “master regulator” of female post-mating responses, needed to “switch” a female from a virgin into a mated state (Kubli and Bopp 2012), but we do not know the identity of gene regulatory networks or key transcription factors in the female that respond to SP, and whether these networks and transcription factors can also be regulated by other reproductive molecules besides SP. In addition, we do not know whether gene networks that are regulated by SP act independently of each other, or whether they are under the control of a shared molecular regulator. This question is also relevant in the context of sexual conflict. While many of SP’s effects increase short-term reproductive output and likely benefit both sexes, some of SP’s effects might be costly to females (for an in-depth review see Hopkins and Perry 2022). For example, SP’s reduction of a female’s remating rate limits the pool of genetic variation she has at hand to pass on to her offspring. Further, there is evidence that the receipt of SP reduces a female’s lifetime reproductive success (Wigby et al. 2005). If some but not all of SP’s effects are costly to the female, the molecular underpinnings of these responses - and more specifically whether they are regulated by a shared molecular mechanism - could impact whether females are able to adapt to resist some of SP’s costly effects without losing access to SP’s beneficial effects.

To address these gaps in our knowledge, we investigated the genetic architecture of the female’s response to SP using time series gene expression data. Fine-scale temporal transcriptomics analysis is a powerful tool to investigate the genetic architecture of a response, as it can distinguish primary from secondary (and later) responses, pinpoint potential regulators and co-regulated targets, and can be used to construct gene interaction networks (e.g. Schlamp et al. 2021; Ciofani et al. 2012; Jo et al. 2021). We measured gene- and exon-level expression changes in the female head at 10 time points within the first 24 hours after mating with a control or a SP null male. We focused on the head to include neuronal, metabolic and endocrine tissues. Our results demonstrate that SP is dispensable for the earliest transcriptomic virgin-to-mated switch, but that it is needed to maintain and diversify post-mating transcriptome changes. We further found that gene regulatory networks that likely underlie specific post-mating responses induced by SP, might be under shared control of the female’s circadian clock.

## Results

To improve our understanding of the genetic architecture of the female’s response to SP, we sampled heads of females that were virgin (V), and at 30 minutes, and 1, 2, 3, 4, 5, 6, 8, 12 and 24 hours after mating with either a control male (SP+) or a SP null male (SP-) (**Fig. 1A**). We sampled more densely at early time points to capture the chronology of the earliest expression changes, and potentially the earliest regulators that act downstream of SP. All matings were set up in the morning and samples of different treatments were collected in parallel to avoid confounding effects caused by the time of day. Below, we describe the results of a differential gene expression analysis and a differential exon use analysis. We then discuss the time frame within which differential expression occurs in the absence or presence of SP. Finally, using a cluster analysis and transcription factor binding motif enrichment analysis, we further explore the biological processes and predicted regulatory networks that are influenced by SP (**Fig. 1B**).

**Figure 1:**
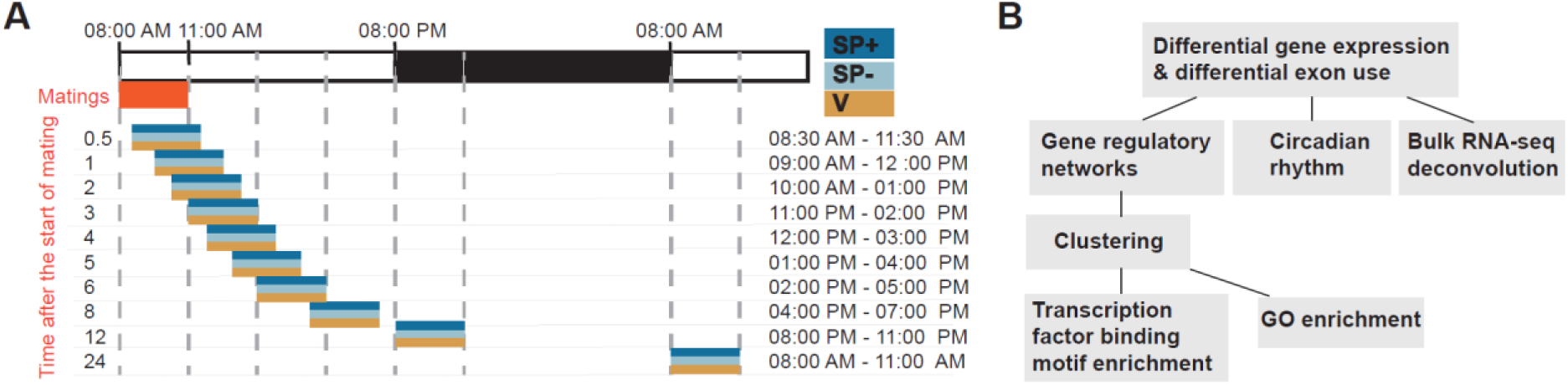
A time series RNA-sequencing dataset of the female head’s response to Sex Peptide and mating. A) Design of the time series mating assay. Three replicates of the time series were run on female heads of virgins (V), females mated to wild-type males (SP+) and females mates to Sex Peptide null males (SP-). B) Overview of the analysis steps.

### Mating and Sex Peptide alter RNA abundance at the gene and exon level

We first performed analyses to detect differential gene expression and differential exon use in response to SP and mating. A differential exon use analysis compares read counts for individual exons across treatments, while controlling for the total read counts across the gene body (see Materials & Methods). Changes in exon usage suggest that alternative splicing might occur or, if the most 5’ exon is differentially used, the differential exon use could indicate a change in preferred transcription start site after mating, perhaps driving expression changes in specific tissues or cell types. In the context of mating, two studies have demonstrated a role for *Homothorax* and *Eip75-B* splice isoforms in female post-mating responses (Garaulet et al. 2021; Zipper et al. 2020). A genome-wide study was done by (Diaz et al. 2021), who quantified alternative exon use in the heads of mated *D. mojavensis* and *D. arizonae* females. However, genome-wide alternative transcript isoform usage has so far received little attention in the context of mating in *D. melanogaster*, and nothing is known regarding its response to SP.

Using pairwise comparisons between the three treatments, with a *q-value* threshold of 0.05 and requiring at least a 1.5-fold change in expression at minimum one time point, we found 666 genes and 86 exons with differential read counts across treatments **(Table S1, S2)**. Of these differentially expressed features, 223 genes and 20 exons underwent at least one 2-fold change in expression between the treatments. The 86 exons mapped to a total of 74 genes. We found little overlap between genes that are differentially expressed across the gene body, and genes that contain one or more differentially used exons: only 12 genes were found in both categories (**Fig. 2A**). Genes that are differentially expressed across the gene body, and genes with differential exon use, were enriched for distinct GO terms (**Fig. 2B, C**). These results are in line with results from Diaz et al. (2021). Similarly to our results, Diaz et al. (2021) report a “functional specialization” of genes that are differentially expressed at the gene versus exon level. Diaz et al. (2021) report that genes with alternative exon use were among others enriched for “muscle assembly” functions, and that differentially expressed genes were enriched for metabolic functions. We likewise found that genes with differential exon use are among others enriched for “regulation of actomyosin structure” (**Fig. 2B**), and that differentially expressed genes have metabolic roles (**Fig. 2C**). This suggests that some of these post-mating responses might be conserved, and that similar strategies for gene regulation are employed. Taken together, mating and SP alter RNA abundance at both the gene- and individual exon-level, and analyses at both levels yield complementary information.

**Figure 2:**
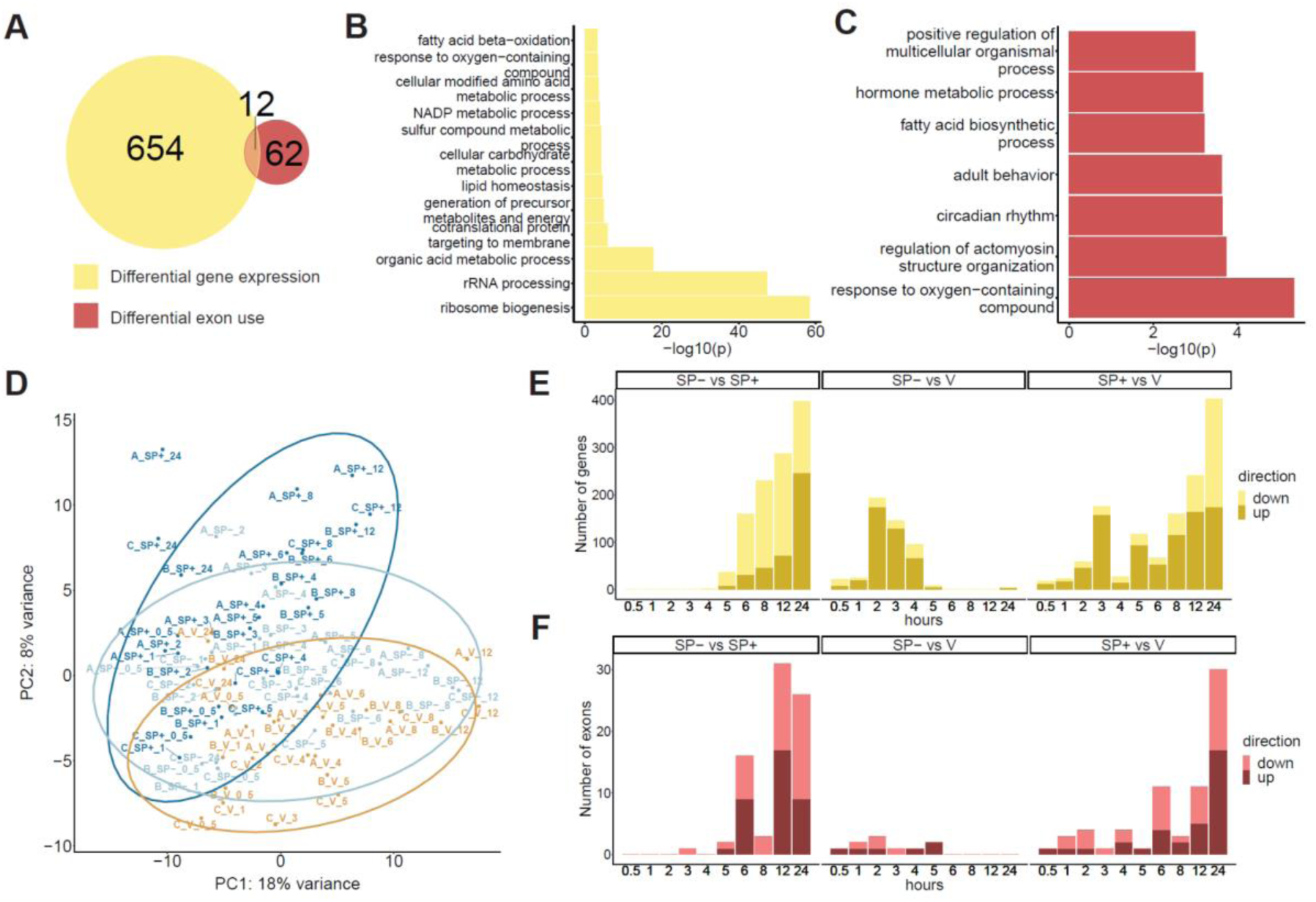
Mating and Sex Peptide alter RNA abundance at the level of genes and individual exons, and act within different timeframes. A) Number of differentially expressed genes, and number of genes with significant differential exon use at different time intervals after mating, with or without SP. B) Significantly enriched (*q*-value < 0.05) GO terms for differentially expressed genes. C) Significantly enriched (*q*-value < 0.05) GO terms for genes with differential exon use. D) Principal component plot generated using gene-level normalized counts for the 500 most variable genes in the dataset. Samples in the plot are indicated using replicate (A,B,C), treatment (V = virgin, SP+ = mated with WT male, SP- = mated with SP null male), and time point. E) Number of differentially expressed genes at each time point, for each pairwise contrast. (*q-value* < 0.05 at the indicated time point; each gene has at least a 1.5-fold change in expression at minimum one of the time points.) F) Number of differentially used exons at each time point, for each pairwise contrast. (*q-value* < 0.05 at the indicated time point; each exon has at least a 1.5-fold change in expression at minimum one of the time points.)

### Sex Peptide is dispensable for the earliest post-mating transcriptome changes

To understand the temporal dynamics of expression changes after mating and in response to SP, we performed a principal component analysis. This analysis showed that mated SP-samples cluster with SP+ samples at early time points, up to ∼ 2-4 hours after mating (along PC2). At later time points, SP-samples group with V samples (along PC1) (**Fig. 2D**). We saw a similar pattern when plotting the number of significant expression changes at each time point for each pairwise contrast, and strikingly similar results were found for differentially expressed genes and differentially used exons (**Fig. 2E, F**). When contrasting SP+ and V, we observed two waves of transcriptome changes: a smaller one before 4 hours and a larger one after 4 hours post-mating. The first, smaller wave of transcriptome changes is SP-independent, because it is also seen in SP- vs. V. The second, larger wave of transcriptome changes depends on SP, since it is only present in SP+ vs V. Thus, non-SP aspects of mating trigger short-term transcriptome changes, but SP is essential for long-term changes, starting from around 5 hours after mating. This timing coincides with the time at which sperm storage is often completed and unstored sperm and seminal fluid proteins have been ejected (Lee et al. 2015), leaving Sex Peptide, bound to stored sperm, present as one of the main remaining seminal fluid proteins in the female. The early SP-independent transcriptome changes are also in line with earlier reports that showed that non-seminal fluid protein components of mating induced responses in mated females within the first 3 hours after mating (McGraw et al. 2004; Heifetz et al. 2014; Tram and Wolfner 1998).

### A clustering approach to construct gene regulatory networks altered by mating and Sex Peptide

To identify gene regulatory networks that are influenced by mating and SP, we grouped genes and exons with similar expression profiles. For this analysis, after removing a small set of outliers (see Materials & Methods), we used 83 differentially used exons and examined a larger set of 1,222 differentially expressed genes (**Table S1**). These genes were selected using a more lenient minimum expression difference of 30% (based on shrunken log_2_ fold change estimates), but still with a *q-value* < 0.05. Because of the patterns in the principal component analysis, we assigned these 1,305 features (genes and exons) to one of three groups. The first group contained 368 features whose first significant change in expression was induced by a non-SP aspect of mating, and occurred within 30 minutes to 4 hours post-mating. The second group contained 369 features whose first significant change in expression was induced by SP, and occurred within 5 to 12 hours post-mating. The final group contained 568 features whose first significant change in expression was induced by SP, at 24 hours post-mating. (**Fig. 3**). Next, we applied hierarchical clustering using lead-lag correlation (LLR^2^) values as a similarity measure (Venkatraman et al. 2021; see Materials and Methods), to identify smaller clusters of features within each of the three groups. We identified 25 clusters within group 1, 16 clusters within group 2, and 21 clusters within group 3. On each of these clusters, we ran a motif enrichment (Aibar et al. 2017) and GO enrichment analysis, and we made use of publicly available single-cell RNA-seq data of the female head to predict in which cell types these expression changes might occur (Sokolowski et al. 2021; Li et al. 2021).

**Fig. 3:**
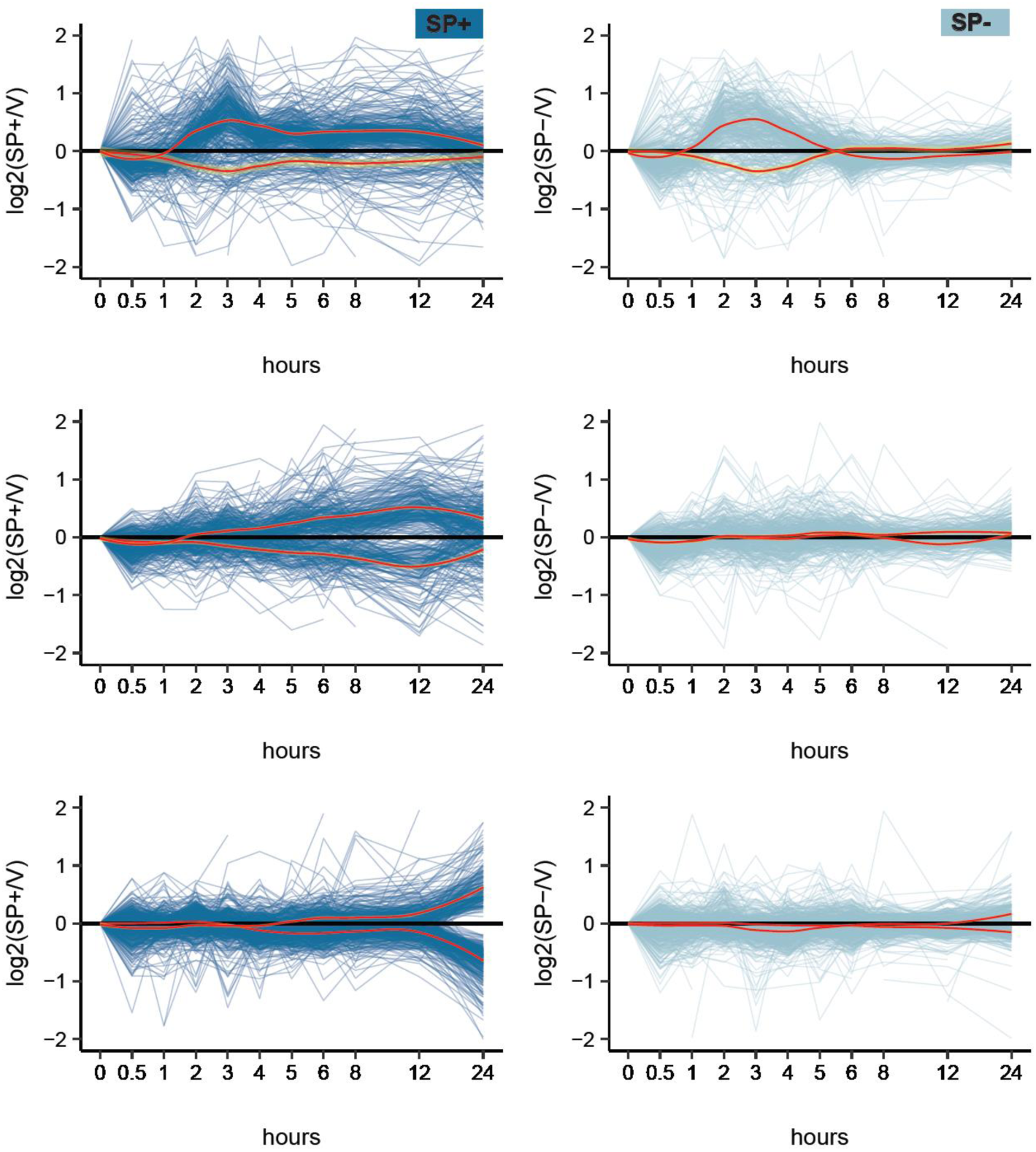
Temporal expression changes induced by mating and Sex Peptide. Y-axis limits were set to -2 and +2 to show representative expression changes more clearly. Red trendlines were obtained using lowess smoothing. Yellow shaded areas show 95% confidence intervals. Within each group, lowess smoothing was applied separately to genes with a mostly up- or down-wards trend in SP+ females (for group 1 based on up- versus down-regulation at 3 hours post-mating; for group 2 based on up- versus down-regulation at 12 hours post-mating; for group 3 based on up- versus down-regulation at 24 hours post-mating).

### A non-Sex Peptide aspect of mating upregulates a gene network with roles in cell growth and protein biosynthesis, and Sex Peptide is needed to sustain these changes

Among the 368 features that undergo significant, SP-independent changes in expression within 30 minutes to 4 hours post-mating, we identified 17 genes that encode transcription factors (*CG4360, E(spl)mbeta-HLH, E(spl)m3-HLH, Clk, Kr-h1, cbt, Myc, cwo, Dr, unc-4, dsx, CrebA, sr, Hr38, Eip74EF:E005, h, tj*), suggesting that non-SP signals from mating can alter transcriptional regulatory networks in the female head. While SP is not required for these earliest mating-induced changes, SP is essential to maintain differential expression of these features between 5 and 24 hours post-mating (**Fig. 3, S11**).

Using a motif enrichment analysis, we identified four transcriptional activators (*CrebA, Clk, Myc* and *tj*) and one transcriptional repressor (*Blimp-1*) that are predicted to regulate features in this group. All four transcriptional activators are themselves among the 368 features, while the transcriptional repressor is significantly differentially expressed in our dataset, but with a < 30% difference between mated and virgin females (thus it was initially not selected for our cluster analysis). In most cases, the transcriptional regulators undergo expression changes 1 to 2 hours before expression changes in their targets are visible (**Fig. 4B-D, B’-D’**).

**Figure 4:**
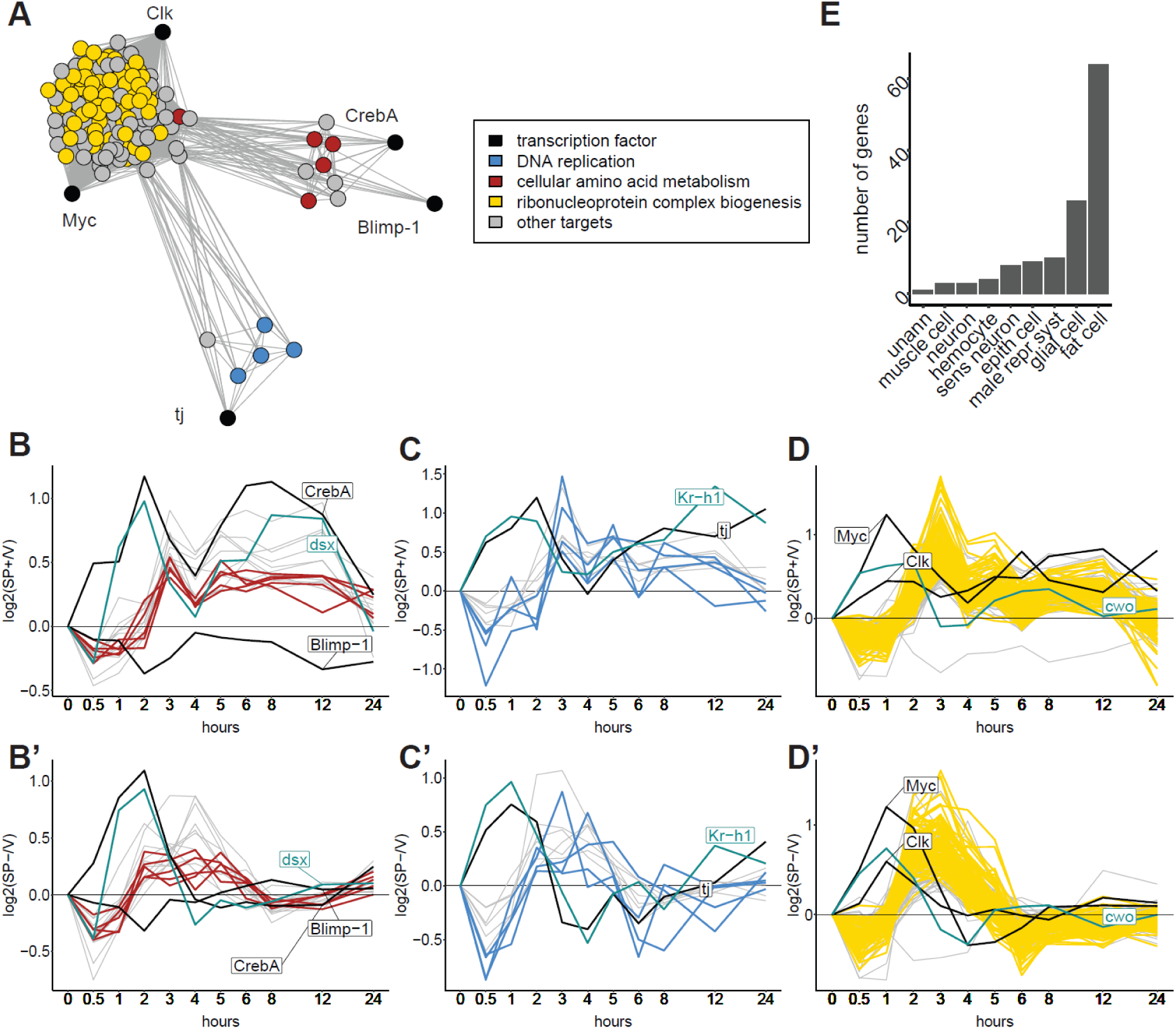
Within 4 hours after mating, expression changes are seen in features with roles in cell growth and protein biosynthesis. A) Network showing transcription factors and predicted target genes, constructed using temporal expression differences between SP+ and virgin females. Edges represent distances between features based on LLR^2^ values. Only edges are shown between transcription factors and predicted targets, and among predicted targets of the same transcription factor. Nodes are colored based on significant (*q*-value < 0.05) GO term enrichment. B-C-D) Time series log_2_ fold changes of mated (SP+) versus virgin (V) females for transcription factors and predicted targets. Transcription factors with similar expression patterns are highlighted in green. B’-C’-D’) Time series log_2_ fold changes of mated (SP-) versus virgin (V) females for transcription factors and predicted targets. E) Assignment of differentially expressed genes to annotated cell types according to scMappR analysis (sens neuron = sensory neuron; epith cell = epithelial cell; unann = unannotated; male repr syst = male reproductive system).

First, we identified a subnetwork containing *CrebA* (upregulated after mating) and the transcriptional repressor *Blimp-1* (downregulated after mating), whose targets are enriched for “cellular amino acid metabolism” (**Fig. 4A, B, B’**). *CrebA* is a regulator of genes involved in the canonical secretory pathway (Johnson et al. 2020). *Blimp-1* plays a role in the fat body during pupal development (Akagi et al. 2016). Motif enrichment analysis did not identify targets for the transcription factor *dsx*, but *dsx*’s expression pattern in response to mating and SP is highly similar to that of *CrebA* (**Fig. 4B, B’**). *dsx* is known to be upregulated in the fat body after mating, where it can bind directly to an enhancer that stimulates the expression of yolk proteins (Burtis et al. 1991; An and Wensink 1995), which are secreted into the hemolymph to reach the ovary. Second, we identified a subnetwork containing the transcription factor *tj*, best studied for its role in gonad morphogenesis (Panchal et al. 2017), as a potential regulator of genes with roles in “DNA replication” (**Fig. 4A, C, C’**). We did not identify predicted target genes for the differentially expressed transcription factor *Kr-h1*, but its expression pattern is similar to that of *tj* (**Fig. 4C, C’**)*. Kr-h1* is part of the flies’ response to Juvenile Hormone (Minakuchi et al. 2008) (whose titers are upregulated by SP (Moshitzky et al. 1996; Bontonou et al. 2015)), and *Kr-h1* is needed for mating-dependent gut remodeling (Reiff et al. 2015). A final subnetwork contains the transcription factors *Clk* and *Myc,* whose predicted targets are enriched for “ribonucleoprotein complex biogenesis” (**Fig. 4A, D, D’**). *Clk* encodes a regulator of the circadian clock (Darlington et al. 1998), and we find that *cwo*, another regulator of the circadian clock, has a similar expression pattern (**Fig. 4D, D’**). The conserved proto-oncogene *Myc* is known to stimulate ribosome biogenesis and cell growth (Grewal et al. 2005). The process of ribosome biogenesis has also been shown to be under circadian control (Abruzzi et al. 2011). Overall, these results suggest that a non-SP aspect of mating stimulates cells in the head to increase their capacity to synthesize proteins, and that SP is required to maintain these changes long-term. By deconvolution of our bulk RNA-seq dataset using publicly available single-cell RNA-seq data of the head, we find that, of the differentially expressed genes that can be assigned to a specific cell type, most are assigned to fat cells (**Fig. 4E**). This suggests that the networks that stimulate cell growth and protein synthesis might be situated in the fat body - an important metabolic tissue and site for yolk protein synthesis and secretion after mating.

### Within 5 to 12 hours post-mating, Sex Peptide sustains increased protein biosynthesis and alters carbohydrate metabolism

We found 369 features that undergo their first significant, SP-induced change in expression within 5 to 12 hours after mating. Most of these features undergo their maximal change in expression between mated (SP+) and virgin females at 12 hours after mating, which is nighttime in our time series. This set of features includes 12 transcription factors and two exons of genes that encode transcription factors (*Pdp1:E014, dsx:E007, Hr4, dl, Lime, Stat92E, ssp, CTCF, CG11695, CG12744, CG14655, CG30431, CG32121, CG3847*). Using a motif enrichment analysis, we found 8 transcription factors that are predicted to regulate genes in this group (**Fig. 5A**). These include transcription factors that are themselves part of this group, and ones that were part of group 1, i.e. underwent mating-induced changes that occurred within the first 4 hours post-mating.

**Figure 5:**
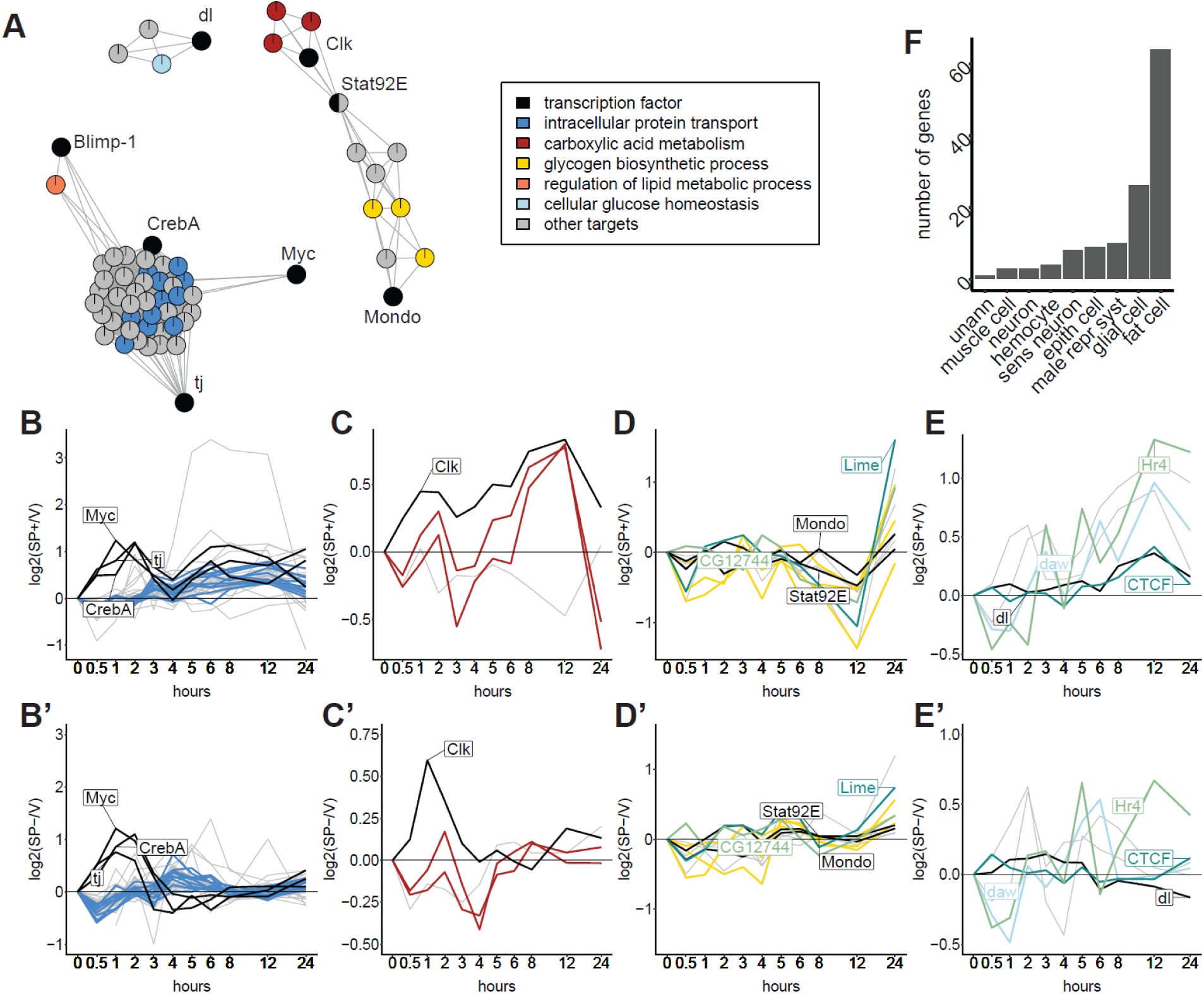
Within 5 to 12 hours after mating, Sex Peptide sustains increased protein biosynthesis and alters carbohydrate metabolism. A) Network showing transcription factors and predicted target genes, constructed using temporal expression differences between SP+ and virgin females. Edges represent distances between features based on LLR^2^ values. Only edges are shown between transcription factors and predicted targets, and among predicted targets of the same transcription factor. Nodes are colored based on significant (*q*-value < 0.05) GO term enrichment. B, C, D, E) Time series log_2_ fold changes of mated (SP+) versus virgin (V) females for transcription factors and predicted targets. Transcription factors with similar expression patterns are highlighted in green. B’, C’, D’, E’) Time series log_2_ fold changes of mated (SP-) versus virgin (V) females for transcription factors and predicted targets. F) Assignment of differentially expressed genes to annotated cell types according to scMappR analysis (sens neuron = sensory neuron; epith cell = epithelial cell; unann = unannotated; male repr cyst = male reproductive system).

First, *Myc*, *CrebA* and *tj* are predicted to upregulate genes involved in “intracellular protein transport” (**Fig. 5A, B, B’**). We found that *dsx:E007* has a similar expression pattern to these transcription factors. *dsx:E007* is an exon of the *dsx* transcription factor, and its RNA abundance has a similar profile over time as that of the entire *dsx* gene body (**Fig. S12**). *dsx:E007* is part of the most 5’ exon that is unique to *dsx* isoforms *RC* and *RD*, suggesting that at least one of these is transcribed more than other *dsx* isoforms in response to SP. Among the genes that respond to SP within 5 to 12 hours after mating, we also found *Yp2* (*Yolk protein 2*), a known *dsx* target (Burtis et al. 1991; An and Wensink 1995) (**Fig. S12**). Second, among the predicted targets of the transcription factor *dl* (*dorsal*) is the gene *daw,* which is involved in glucose homeostasis (Chng et al. 2014) (**Fig. 5A, E, E’**). Also upregulated are the transcription factor *Clock* and most of its predicted targets, which are involved in carboxylic acid metabolism (**Fig. 5A, C, C’**). Additional upregulated transcription factors for which no predicted targets were found using motif enrichment analysis, include *CTCF* and *Hr4*, which are both involved in the response to ecdysone (Pascual-Garcia et al. 2017; King-Jones et al. 2005), a hormone that is upregulated after mating (Harshman et al. 1999) (**Fig. 5E, E’**). We also found one upregulated gene that is involved in the regulation of lipid metabolism, and which is a predicted target of the transcriptional repressor *Blimp-1* (**Fig. 5A**). Finally, the transcription factors *Stat92E* and *Mondo* are predicted to regulate genes involved in glycogen biosynthesis. These transcription factors and their targets are downregulated at 12 hours post-mating (**Fig. 5A, D, D’**). *Mondo* is a known regulator of lipid and carbohydrate metabolism, and is itself regulated by glucose availability (Musselman et al. 2013; Chatterjee and Perrimon 2021). Additional transcription factors with no predicted targets but with similar expression patterns to *Mondo* and *Stat92E* are *Lime* and *CG12744* (**Fig. 5D, D’**)*. Lime* (*Linking immunity and metabolism*) was previously found to be crucial for energy provisions during an immune response, due to its role in maintaining levels of glycogen and trehalose (Mihajlovic et al. 2019). Deconvolution of our bulk RNA-seq data at 5 to 12 hours post-mating indicates that most differentially expressed genes that can be assigned to a specific cell type are assigned to “fat cell” (**Fig. 5F**). Taken together, at 5 to 12 hours post-mating, SP continues to stimulate responses related to protein biosynthesis, which were already initiated in the first 4 hours after mating by a non-SP aspect of mating, and which likely occur in the fat body. In addition, SP alters the expression of features involved in carbohydrate metabolism, which has been described before (White et al. 2021), but we observe that the most significant changes occur at night.

### 24 hours after mating, Sex Peptide downregulates a gene network with neuronal functions

We identified 568 features that undergo their first significant change in expression 24 hours after mating, which is the next morning in our time series. These responses are entirely SP-specific. Across all 568 features in this group, there is a highly significant enrichment of genes involved in “generation of energy and precursor metabolites” and “cytoplasmic translation” (respective *q-values* = 5.7 x 10^-8^ and 5.1 x 10^-6^), and almost all of these features are downregulated in a SP-dependent manner (**Fig. 6B**). Similar GO terms were also found to be downregulated at 3 hours post-mating in head-thorax samples by Fowler et al. (2019).

**Figure 6:**
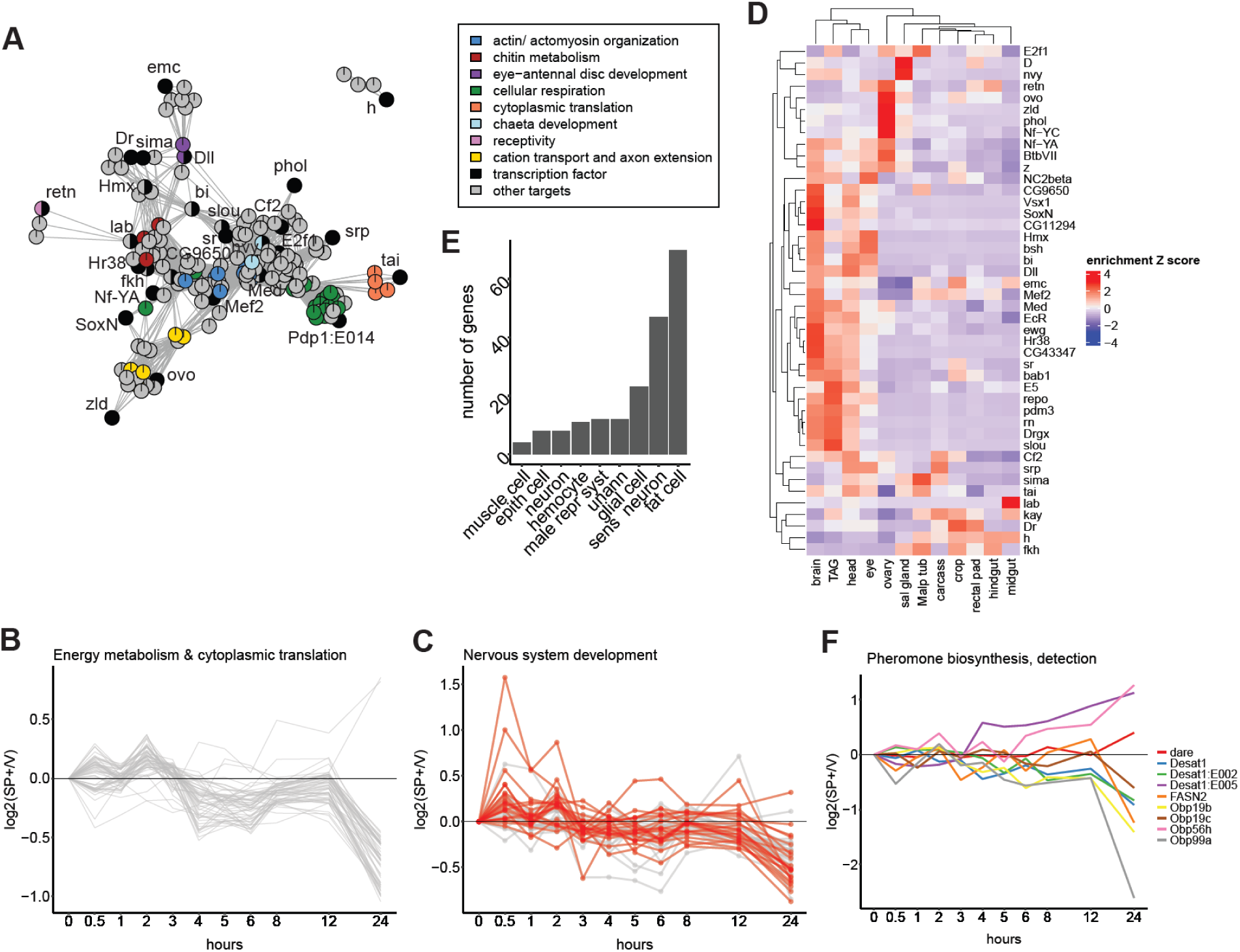
After 24 hours, Sex Peptide downregulates genes with roles in the nervous system. A) Network showing transcription factors and predicted target genes, constructed using temporal expression differences between SP+ and virgin females. Edges represent distances between features based on LLR^2^ values. Only edges are shown between transcription factors and predicted targets, and among predicted targets of the same transcription factor. Nodes are colored based on significant GO term enrichment. B) Time series log_2_ fold changes of mated (SP+) versus virgin (V) females for genes involved in cytoplasmic translation and energy generation. C) Time series log_2_ fold changes of mated (SP+) versus virgin (V) females for transcription factors with maximal expression changes at 24 hours after mating, and transcription factors predicted to regulate targets that have maximal expression changes at 24 hours post-mating. Transcription factors associated with the enriched GO term “nervous system development” are highlighted in red. D) Tissue-specific expression normalized to whole body expression, based on FlyAtlas 2 (TAG = thoracic-abdominal ganglion, Malp tub = Malpighian tubule, sal gland = salivary gland). E) Assignment of differentially expressed genes to cell types by scMappR (sens neuron = sensory neuron; epith cell = epithelial cell; unann = unannotated; male repr cyst = male reproductive system). F) Time series log_2_ fold changes of mated (SP+) versus virgin (V) females for genes with roles in pheromone biosynthesis and detection.

A motif enrichment analysis identified 27 transcription factors predicted to regulate genes in this group. These transcription factors and most of their predicted targets were downregulated in response to SP. Predicted targets are enriched for “actin cytoskeleton organization” and “actomyosin structure organization”, “cellular respiration” and “cytoplasmic translation”, “chitin metabolism”, antennal and chaetal development, “axon extension” and “cation transport”, and the “regulation of female receptivity” (**Fig. 6A**). In addition to the 27 transcription factors identified using motif enrichment analysis, there are another 18 transcription factors present in this group, for which no predicted targets were found. Among this total set of 45 transcription factors, there is a significant enrichment of the GO term “nervous system development” (23 genes, **Fig. 6C**), and 18/45 have enriched expression (relative to whole body expression) in the fly brain or thoracic-abdominal ganglion, based on FlyAtlas 2 data (Leader et al. 2018); **Fig. 6D**). The majority of the 45 transcription factors are downregulated in a SP-dependent manner, with the strongest downregulation at 24 hours.

In addition to genes with neuronal functions, this group also contains genes with roles in pheromone biosynthesis and detection (*dare, Desat1, FASN2, Obp19b, Obp19c, Obp56h, Obp99a*, **Fig. 6F**) (Freeman et al. 1999; Chung et al. 2014; Bousquet et al. 2012; Xiao et al. 2019). For *Desat1*, a desaturase involved in cuticular hydrocarbon and pheromone metabolism (Wicker-Thomas et al. 1997; Marcillac et al. 2005), we further detected two differentially used exons (*E002* and *E005*), which make up two of a total of five alternative first exons of the *Desat1* gene (Bousquet et al. 2012). Differential exon use started already within the first 12 hours post-mating, and the exons show expression changes in opposite directions (**Fig. 6F**). Deconvolution of the bulk RNA-seq data at 24 hours assigned the largest number of genes (63 genes) to “fat cell”, but in second place is “sensory neuron” with 43 genes (**Fig. 6E**).

Together, the enriched GO terms, gene functions and tissue- and cell type-enrichment suggest that, while earlier responses mostly affected protein and carbohydrate metabolic processes and potentially took place in the fat body, at least a subset of these late responses are more likely to take place in neuronal tissues in the head. We speculate that the downregulation of neuronal transcription factors and predicted targets might reflect a decreased function of certain neurons in the mated female’s head. This could also explain the coordinated downregulation of genes involved in translation and respiration, since energy metabolism is often used as a reflection of a neuron’s activity (Thompson et al. 2003). A decreased activity of certain neurons in the head could contribute to altered sensory and behavioral responses in mated females that received SP.

### Mating and Sex Peptide alter the expression of features that have a circadian rhythm in virgin females

Because we found that a non-SP aspect of mating (in the first 4 hours post-mating) and SP (after 4 hours) upregulated the expression of *Clock,* we investigated whether additional features with circadian rhythms were differentially expressed after mating. Using the R package MetaCycle (Wu et al. 2016), we found that of the 1,305 differentially expressed genes and differentially used exons, 18% (232 features) follow a circadian rhythm in virgin females (*q-value* < 0.3, **Fig. 7A**). These include known regulators of the clock (*Clk*, *cwo, tim, CG31324/Cipc* and an exon of *Pdp1*, *Pdp1:E014;* **Fig. 7D, 7C**), and two genes that were found to be putative regulators of circadian rhythms by Litovchenko et al. (2021) (*CG14688* and *Gclm*).

**Figure 7:**
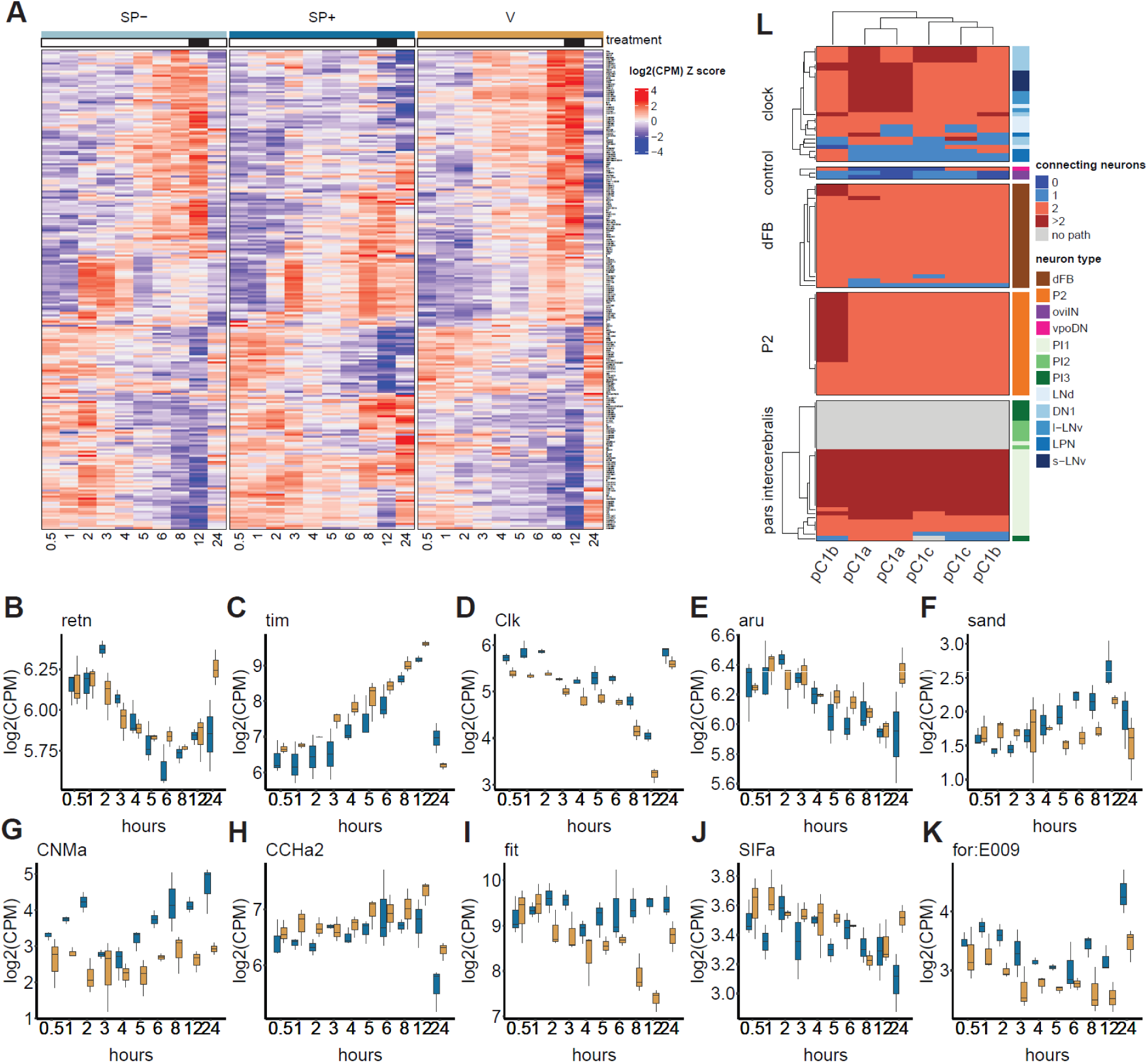
Mating and Sex Peptide alter the expression of genes and exons that follow a circadian rhythm in virgin females. A) Row-normalized Z scores of median log_2_(CPM) (counts per million) across available biological replicates, for genes that have a circadian rhythm in virgin females according to MetaCycle analysis. B-K) log_2_(CPM) values for selected genes. L) pC1 neurons (pre-synaptic) have (in)direct connections to neurons that regulate circadian behaviors (post-synaptic), based on the Fly Hemibrain connectome (v1.2.1; Scheffer et al. 2020). dFB = dorsal fan-shaped body; control = oviIN and vpoDN neurons, which are known to be involved in the female post-mating response downstream of pC1 neurons; clock = central clock neurons.

For the 368 features that already show expression changes within the first 4 hours after mating, induced by a non-SP aspect of mating, 73 (20%) have a circadian rhythm in virgin females. These include genes involved in ribonucleoprotein complex biogenesis, which are upregulated at night in virgin females, but are also upregulated during the day in mated females (**Fig. 4D, D’**). Of the 369 features that show SP-induced expression changes within 5 to 12 hours post-mating, 91 (25%) have a circadian rhythm in virgin females. These include features involved in glycogen biosynthesis and the transcription factor *Lime*, which have lower expression in mated (SP+) vs. virgin females at night (**Fig. 5D, D’**). These results suggest that SP might act on the female’s circadian clock to mediate metabolic changes - in this case to stimulate protein synthesis throughout the day and downregulate glycogen metabolism at night. For the 568 features with significant differential expression at 24 hours after mating, 68 features (12%) have a circadian rhythm in virgin females. Among these are *aru,* which plays a role in ethanol sensitivity and synapse number (Eddison et al. 2011) and *retn*, which has been implicated in regulating female receptivity to male courtship (Ditch et al. 2005; Terhzaz et al. 2007). These genes are downregulated in the evening and at night in virgin females and upregulated in the morning. In mated females that received SP, *aru* and *retn* remain downregulated the next morning (**Fig. 7B, E**). Female mating activity is a circadian behavior, and females are most receptive in the morning (Sakai and Ishida 2001). Aside from receptivity, other SP-induced post-mating responses affect circadian behaviors, such as increased feeding (Uchizono et al. 2017; Carvalho et al. 2006), and reduced siesta sleep (Isaac et al. 2010). Among our differentially expressed features, we found six candidates that might act downstream of SP and the circadian clock to influence feeding (*fit* (**Fig. 7I**; Sun et al. 2017)*, CNMa* (**Fig. 7G**; Kim et al. 2021)*, CCHa2* (**Fig. 7H**; Ren et al. 2015*), SIFa* (**Fig. 7J**; Martelli et al. 2017) and *for:E009*) and activity (*CNMa* and *sand*). We note that *CNMa* and *sand* were not among the genes with a circadian rhythm in virgin females in our dataset based on the MetaCycle analysis. However, *sand* is involved in sleep regulation (Pimentel et al. 2016), and *CNMa* is known to be expressed in clock neurons (Jin et al. 2021; Ma et al. 2021). One of these six candidates is an exon (*E009*) of the *foraging* gene. This exon is upregulated after mating and in response to SP. In virgin females, counts of *for:E009* go down during the day (**Fig. 7K**). Exon *009* is part of a subset of transcripts of *foraging*, which are all under the control of the fourth promoter of the *foraging* gene. We used quantitative PCR to confirm that indeed only transcripts of *foraging*’s 4th promoter are upregulated after mating, and not transcripts controlled by promoters 1, 2 or 3 (**Fig. S13**). Anreiter et al. (2017) have shown that transcripts expressed from *foraging*’s fourth promoter influence foraging behavior in adult females, but a difference in foraging behavior between mated and virgin females was not studied. Another candidate, *sand* is upregulated in SP+ versus virgin females at 5 to 12 hours post-mating (**Fig. 7F**). *sand* encodes a potassium channel, and its function has been described in neurons of the dorsal fan-shaped body in the brain, which are involved in sleep homeostasis. sand protein mediates potassium leakage from these neurons, thereby contributing to keeping them in a quiescent state that is typically found in well-rested flies (Pimentel et al. 2016). The timing of *sand* upregulation in our dataset coincides with the afternoon, which is also the time at which mated females are more active than virgin females (Isaac et al. 2010). Together, these results demonstrate that a non-SP aspect of mating, and SP can alter the transcript abundances of circadian genes, several of which have functions related to post-mating responses. These alterations to the circadian rhythm are only maintained past 5 hours post-mating in the presence of SP. Our results further suggest that this could potentially be achieved by SP’s effect on the expression of core molecular components of the clock.

### Sex Peptide-responsive neurons connect to neurons that regulate circadian behaviors in the brain

One possible mechanism for how SP can alter females’ circadian expression patterns, is via connections between neurons known to respond to SP and neurons that regulate circadian behaviors in the brain. To investigate this possibility, we used the neuPRINT+ connectome database’s “Shortest Path” search (Scheffer et al. 2020). We examined connections from neurons that respond to Sex Peptide (pC1) to neurons involved in the regulation of circadian behaviors, including feeding and sleeping (central clock neurons (Patke et al. 2020), P2 neurons (Li et al. 2021), neurons of the Pars Intercerebralis (Cavanaugh et al. 2014) and dorsal fan-shaped body neurons (Donlea et al. 2014)). We examined the most direct links that involved two or fewer connecting neurons. The results of this search did support that there are paths from neurons that respond to SP to neurons involved in regulating circadian behaviors (**Fig. 7L**, **Table S6**). As reported previously, we observed direct connections between pC1 neurons and oviIN neurons (“oviposition inhibitory neurons”, which regulate oviposition) and vpoDN neurons (involved in auditory responses) (Wang et al. 2020, Wang et al. 2021). One other direct connection was found from a pC1b neuron to an LNd clock neuron. pC1 neurons also connected indirectly via one connecting neuron to subsets of LPN, LNd and DN1 neurons of the clock, ExR3 neurons of the dorsal fan-shaped body (ExR3 neurons are involved in regulating sleep-wake cycles (Liu et al. 2019)), and a small number of neurons in the Pars Intercerebralis. Other paths consisted of two or more connecting neurons and, in some cases that involved neurons of the Pars Intercerebralis, no connection was found. We further found that SMP454 and oviIN neurons were frequently used to connect pC1 neurons to neurons that regulate circadian behaviors. Of the paths with 1 or 2 connecting neurons, oviIN neurons were used in the path in 43% of cases, and SMP454 neurons in 52% of cases (**Table S6**), suggesting that these neurons might be significant in transmitting signals downstream of SP. This observation also suggests that oviIN neurons might have a more general role in relaying SP signaling, not only targeted to oviposition regulation, but more generally allowing SP to alter signaling in neurons that regulate circadian behaviors.

## Discussion

Reproduction is only possible by successfully completing a set of defined temporal events. In *Drosophila melanogaster,* for example, mating does not take place without prior courtship, which consists of a series of complex stereotypical, timed behaviors, including tapping, licking, and singing (Bastock and Manning, 1955). Only if a female deems a male suitable will copulation occur. This act is followed by another series of defined, timed events, which include sperm storage, followed by the ejection of unstored sperm and seminal fluid proteins, and the start of increased ovulation (Lee et al. 2015; Herndon and Wolfner 1995). Here, we show that stereotypical, timed events also occur in an additional aspect of mating, namely in the female head’s transcriptomic post-mating response. This transcriptomic response includes both gene-level changes and changes in exon usage, which point to the use of alternative transcript isoforms after mating, an under-studied aspect of the post-mating response in *D. melanogaster*. We found that metabolic changes pertaining to protein biosynthesis were among the first to occur within 4 hours after mating, followed by changes in carbohydrate metabolism after 8-12 hours. Changes in genes with neuronal functions occurred much more slowly, with maximal differences seen only 24 hours after mating. We further found that transcriptome changes accumulated over time, with the largest number of differentially expressed features found at the latest time point sampled. Dalton et al. (2010), who measured gene expression in the mated female’s head up to 72 hours post-mating, also reported peak differential expression at their latest time point. These dynamics appear different in the female reproductive tract, where studies consistently report observing the largest number of expression changes at earlier rather than later time points post-mating (Mack et al. 2006; Prokupek et al. 2009; McDonough-Goldstein et al. 2021). Our observations suggest that different tissues undergo distinctive and timed post-mating transcriptome changes, and highlight the value of studying these changes using fine-scale temporal transcriptomics.

In addition to investigating the effect of mating, we incorporated the effects of the seminal fluid protein Sex Peptide into our time series. Our results highlight that SP is not needed for the initial virgin-to-mated switch, which involves mostly metabolic features. However, SP is needed to initiate a second wave of transcriptome changes, which consists of the maintenance of early metabolic changes and the induction of additional changes in both metabolic and neuronal features. The end of the first wave and start of the second wave coincide with the timing at which unstored sperm and seminal fluid proteins are typically ejected by the female. Thus, seminal fluid proteins that do not remain in the female long-term could act redundantly with SP to induce the expression changes in the first wave. One candidate is the seminal fluid protein Dup99B. Dup99B has a high sequence similarity to SP (Saudan et al. 2002) and can bind the Sex Peptide Receptor (Yapici et al. 2008), but is not found on sperm for longer than 5 hours after mating (Peng et al. 2005; Kubli 2003). However, other seminal fluid proteins, sperm and non-sperm non-seminal fluid protein components of mating, such as pheromones or mechanical cues from copulation, might also be involved in short-term gene expression changes (McGraw et al. 2008; McGraw et al. 2004; Shao et al. 2019; Tram and Wolfner 1998). We further note that a recent report showed that males that lack SP demonstrate altered transfer of a subset of seminal fluid proteins (Wainwright et al. 2021). This altered seminal fluid protein composition could also contribute to the patterns we observe.

Similar to the two waves of transcriptome changes that we identify, McGraw et al. (2008) described two phases in gene expression dynamics in whole females: at 1 to 3 hours post-mating, many small-magnitude changes occur, and at 6 to 8 hours post mating, few large-magnitude changes occur. A two-phase response is also seen regarding changes in female attractiveness within 24 hours post-mating. A mated female’s reduced attractiveness is initially caused by pheromones, but later depends on sperm and accessory gland proteins (a subset of seminal fluid proteins that include SP) (Laturney and Billeter 2016; Tram and Wolfner 1998; Scott 1986). Thus, this two-phase response seems to be a stereotypical pattern that occurs within the first 24 hours after mating in *D. melanogaster*.

The metabolic and neuronal networks we uncovered seemed unrelated at first sight, both in terms of genes and temporal dynamics. However, across both networks we identified genes whose expression follows a circadian rhythm in virgin females. This led us to hypothesize that SP might act on the female’s circadian clock to enable its pleiotropic effects. This hypothesis makes sense in light of our knowledge of post-mating responses, since many of these affect circadian processes like sleep, feeding, receptivity, oogenesis and egg laying (Carvalho et al. 2006; Isaac et al. 2010; Sakai and Ishida 2001; Allemand 1976; Zhang et al. 2021). Because we sampled only within the first 24 hours post-mating, we do not know which responses are recurrent in the next days and nights. Still, there is strong evidence to support the idea that SP influences the female’s circadian rhythm. First, we observed differential expression of known and predicted molecular regulators of the clock. Second, we found that many post-mating differentially-expressed features have a circadian expression pattern in virgin females. Many of these differentially-expressed features have previously been implicated in mating-induced changes in females. For example, it is well established that mating and SP lead to a metabolic shift in females, stimulating dietary protein intake, down-regulating carbohydrate metabolism, and increasing protein biosynthesis (White et al. 2021; Bradley and Simmons 1997; Gioti et al. 2012; Ribeiro and Dickson 2010; Bownes et al. 1988; Uchizono et al. 2017). Post-mating changes in metabolism have been reported in multiple expression profiling studies (Pasquier and Robichon 2022; Fowler et al. 2019; Dalton et al. 2010; White et al. 2021; Domanitskaya et al. 2007), and were also found in our study. However, our sampling pattern allowed us to place the expression changes of these genes, of differentially used exons, and of the transcription factors that are predicted to orchestrate these changes, within the context of day and night, highlighting marked time-of-day-dependent differences with virgin females.

If SP can alter circadian rhythms in females, the question becomes how it achieves this, and at what level SP signals are integrated in the female’s clock. Or, at what level does SP “override” the clock, since circadian expression patterns are altered despite sustained light/dark cycles? Circadian gene expression patterns are regulated on multiple levels. Flies have central clock neurons in the brain that entrain to environmental light/dark cycles (Dubowy and Seghal, 2017), but there are many neurons downstream of these pacemakers, such as neurons in the dorsal fan-shaped body and the pars intercerebralis (Donlea et al. 2014; Donlea et al. 2018; Kempf et al. 2019; Pimentel et al. 2016). In addition, many peripheral tissues, including the fat body, have their own functional clocks (Patke et al. 2020). The metabolic gene regulatory network that we identified, and expression changes of *fit*, suggest that the fat body clock is altered in response to SP. The fat body clock regulates feeding behavior and glucose metabolism, but synchronization between the fat body clock and the clock in neuronal cells is important, and desynchronization has been shown to negatively impact egg laying (Zheng and Sehgal 2008; Fulgham et al. 2021; Erion et al. 2016; Xu et al. 2011). This, in addition to the expression changes we observed in neuronal genes that follow a circadian expression pattern in virgin females (such as *aru, retn, sand*), indicate that circadian changes in response to SP are not likely to be limited to the fat body. Indeed, connections between pC1 neurons and clock neurons form a possible mechanism for SP to directly modulate the female’s clock, thereby allowing SP to control many different post-mating responses at once. This potential mode of action fits with an experimental evolution study that has shown that sexual conflict might drive males to mostly manipulate the female’s central nervous system (Hollis et al. 2019). If SP truly acts on the female’s circadian clock to establish post-mating responses, this could have implications for the evolution of sexual conflict. If SP were to influence independent regulatory networks to trigger post-mating responses, females might more easily adapt to escape one of its effects, such as the SP-induced reduction in receptivity to other males, which benefits the male a female mated with, but not necessarily the female herself. By acting on the female’s circadian rhythm, SP could control many physiological and behavioral changes via one interconnected neuronal network, making it more difficult for the female to escape some of SP’s costly effects without also losing access to beneficial effects.

In conclusion, using a high-resolution time-course RNA-sequencing dataset of female heads, we have shown that a non-SP aspect of mating acts redundantly with SP to regulate a network of metabolic genes in females early after mating, and that SP is needed to sustain and diversify these changes to additional metabolic and neuronal gene regulatory networks. We identified transcription factors that are regulated downstream of SP, and these are likely involved in regulating post-mating responses that influence a female’s metabolism, behavior, or neuronal activity. We further found that these gene networks themselves might be under shared circadian control. Future research opportunities lie in understanding mechanistically how signals from mating wire into the female’s clock system, how that translates into expression changes in peripheral tissues, and what the significance of this is for the evolution of sexual conflict.

## Acknowledgements

We thank A. Jain for help with sample collection and library prep; the Cornell Genomics Facility for library sequencing; N. Brown, D. Chen, Y. Hafezi and A.-M.Jaksic for feedback regarding the experimental design; S. Misra for running the Western blot; E. Cosgrove for help with cluster analysis; I. Anreiter and M. Sokolowski for advice for *foraging* qPCR experiments. A.G. Clark and M.F. Wolfner were supported by NIH award R01 HD059060. S.Y.N. Delbare, S. Venkatraman, A.G. Clark, M.T. Wells and S. Basu were supported by NIH award R01 GM135926. S. Basu also acknowledges support from NIH award R21NS120227 and NSF award DMS-1812128.

## Data Availability

Fastq files and raw gene- and exon-level count data tables are available on GEO (GSE198879). The R code used for analyses is available on GitHub (SDelb/Transcriptomics_SP_timecourse).

## Materials & Methods

### Fly stocks and husbandry

All flies were maintained on standard yeast/ glucose medium in a 12 hour light/ dark cycle (lights on 8 AM, lights off 8 PM), at 25°C. Females used in the experiment were from a wildtype *Canton-S* strain. Males used in the experiment were offspring of *SP^0^/TM3,Sb* (*SP* null mutant) and *Δ130/TM3,Sb* (a deficiency line with a deletion in the locus containing *SP*) (Liu and Kubli 2003). From this cross, which was set up in both directions, male offspring with wildtype bristles are *SP* null, and male offspring with stubble bristles are controls. These controls are heterozygous for *SP,* but this was shown to be sufficient to induce SP-mediated responses in females (Liu and Kubli 2003). Further, we will refer to virgin females as “V”, females mated to control males as “SP+” and females mated to *SP* null males as “SP-”. All males and females used in experiments were virgin and were aged for three to five days after eclosion.

A Western blot was performed as in Misra and Wolfner (2020) to verify the absence of SP in *SP^0^*/*Δ130* males and the presence in *Δ130/TM3,Sb* and *SP^0^/TM3,Sb* males (**Fig. S1B**). In addition, we performed a four-day receptivity assay to verify the absence of SP in *SP* null males. As expected, within 1 hour, SP- females readily remated with a *Canton-S* male four days after their initial mating with a *SP* null male. This was not the case for SP+ females that had initially mated with a control male (**Fig. S1A**; *n* re-mated for SP- = 13/16; *n* re-mated for SP+ = 4/14; Fisher’s exact test *p-value* = 0.009). In addition to measuring receptivity, we also measured mating duration. We did not detect a significant difference in mating duration between SP null and control males (mean mating duration for SP+ = 25 minutes +/- 4; mean mating duration for SP- = 26 minutes +/- 7 minutes).

### Time series mating assay

All matings were performed at room temperature (21°C). We collected three independent biological replicates (indicated with A, B and C) by repeating the entire experiment in three consecutive weeks. For each repeat experiment, flies were derived from independent parental flies. The afternoon before the day of the mating assay, females were isolated in individual vials. On the day of the mating assay, matings were set up starting at 7:30 AM, by adding one male (either *SP* null or control) to each vial with a female fly. Vials with females designated to remain virgin did not receive a male. Females were given three hours time to mate, between 8 AM and 11 AM (**Fig. 1**). Across all replicates, only two flies mated before 8 AM, and these were excluded from the experiment. All matings were observed at least once every five minutes to record the start time of mating and to ensure that each mating lasted at least 10 minutes. Females were frozen using dry ice at their designated time point after mating (30 minutes, 1, 2, 3, 4, 5, 6, 8, 12, or 24 hours after mating), and virgin females were frozen in parallel, and stored at -80°C. We obtained 90 samples in total (3 treatments x 10 time points x 3 replicates). We used a different method to assign females to a specific freezing time point for replicate A vs. replicates B and C. For replicate A, each female was assigned to a specific time point before the start of the mating assay. For replicates B and C, females were not assigned to a time point ahead of time. Instead, the first female to mate was assigned to be frozen 30 minutes after mating, the second female to mate was assigned to be frozen 1 hour after mating, and so on. Despite the different methods used to assign females to freezing time points, a replicate batch effect was not apparent in a principal component analysis of the RNA-seq counts (see below).

### Head collection, library prep & quantitative PCR

To dislodge fly heads from the thorax, vials with female flies were dipped in liquid nitrogen and vortexed for 3x 5 seconds, dipping vials in liquid nitrogen in between each round of vortexing. Dislodged heads were collected in Trizol, homogenized using a mechanical rotor and plastic pestle, and stored at -80°C. For each sample, 3 to 12 heads were collected, with a median of 6.5 heads. To extract RNA, samples in Trizol were thawed on ice. We extracted RNA and performed an on-column DNase treatment using the Directzol RNA microprep kit (Zymo, CA), according to the manufacturer’s instructions. Sample concentration and purity were determined using Qubit and Nanodrop (Thermo Fisher Scientific, MA). RNA integrity was verified for a randomly selected subset of seven samples using the Fragment Bioanalyzer (Agilent, CA) at the Cornell Genomics Facility. The minimum RQN obtained was 9.4. Poly(A)-selected, stranded libraries were made using the NEBnext Ultra II Directional RNA library prep kit and the Poly(A) mRNA magnetic isolation module (New England Biolabs, MA). Library concentrations were determined using Qubit (Thermo Fisher Scientific, MA) and a Fragment Bioanalyzer (Agilent, CA) was used to inspect library size range and ensure absence of primer dimers. Libraries were sequenced using NextSeq500 (Illumina, CA) at the Cornell Genomics Facility. All 90 samples were sequenced in the same lane, and this was repeated five times to obtain adequate read depth.

For qPCR experiments, cDNA was made using SMARTScribe Reverse Transcriptase (Takara Bio, CA). qPCRs were run using LightCycler 480 SYBR Green I Master (Roche) on a Roche Lightcycler 480 instrument. Primer sequences used are those from Allen et al. (2017) and are listed in **Table S3**. Standard curves were made to ensure primer efficiency was above 90%. qPCR results were analyzed as in Taylor et al. (2019) to obtain normalized expression values, using two housekeeping genes (*Act5C* and *alpha-Tubulin*) as internal controls.

### Read quality control, processing and alignment

Quality control of sequenced libraries was performed using FastQC (version 0.11.8; https://www.bioinformatics.babraham.ac.uk/projects/fastqc/). Sequencing adaptors were removed using Trimmomatic (version 0.39) (Bolger et al. 2014), which was also used to trim low quality bases at the 5’ end of reads (using parameters TRAILING:20 and SLIDINGWINDOW:5:20). Reads with a minimum length of 30 bases were retained and FastQC was run again to ensure removal of sequencing adaptors was successful. Reads were aligned to the *Drosophila melanogaster* genome (dm6 version r6.24) using STAR (version 2.7.5a) (Dobin et al. 2013) and per-gene read counts were determined using HTSeq (version 0.11.2) (Anders et al. 2015). Because we obtained five fastq files for each of the 90 samples, we initially ran the QC, alignment and read counting pipeline on each fastq file separately. We then performed a principal component analysis to check for lane effects (**Fig. S2**). Since no lane effects were observed, we merged the five fastq files for each sample. After merging, the median read count per sample was 22.4 million reads.

### Read count filtering, principal component analysis & correction for batch effects

Raw counts were filtered to keep only genes with at least three counts per million (calculated using edgeR; (Robinson et al. 2010; McCarthy et al. 2012) in at least three samples and genes encoded in mitochondrial DNA were removed. This left 9,027 expressed genes in the dataset. We performed a principal component analysis (PCA) to investigate the dataset for potential outlier samples and batch effects. PCA was done on log_2_-transformed TMM-normalized counts per million for the 500 genes with highest row variance in the dataset. We observed unexpected, random clustering of samples (**Fig. S3A**). To understand what caused this, we investigated the top 100 genes that were driving this pattern. Aside from genes with circadian rhythms, such as *period* and *timeless*, which had regular expression patterns across samples (**Fig. S4A**), the top 100 most variable genes in the dataset included 28 immune response genes, with irregular expression patterns across samples (**Fig. S4B**). We further inspected the expression patterns of these 28 immune genes and found that some samples had much higher expression than others (**Fig. S5A**). The increased expression of immune genes was not specific to any replicate, time point or treatment. We think that differences in immune gene expression may have been caused by variability among fly food batches. To correct for this batch effect caused by an unexpected immune response, we used ComBat-seq, a method developed for batch effect correction in RNA-seq data (Zhang et al. 2020). Specifically, we performed hierarchical clustering on all samples based on Euclidean distance calculated from the samples’ expression for the 28 immune genes. We cut the tree to obtain seven clusters and used the resulting cluster assignments in ComBat-seq. ComBat-seq returns a raw counts matrix, corrected for the immune response batch effect. To ensure the corrected counts were suitable for downstream analysis, we first established that the ComBat-seq correction led to more uniform expression of immune genes (**Fig. S5B**). Second, we investigated the ratio of batch-corrected versus original counts for each gene in the dataset. We found 72 genes with a ratio ≥ 2, indicating a large effect of ComBat-seq count correction. For 54/72 genes, the ratio was ≥ 2 in 9 or fewer samples. A GO enrichment test performed using STRING (version 11) (Szklarczyk et al. 2019) showed that this set was enriched for genes encoding odorant binding proteins. For 18/72 genes, the ratio was ≥ 2 in 10 or more samples. These genes were enriched for immune response genes. Thus, the ComBat-seq correction adjusted more strongly the counts of genes whose gene products are involved in interaction with or response to the environment. ComBat-seq had minimal to no effect on the counts of the remaining 8,955 genes in the dataset. Third, we compared inter-replicate variability before and after Combat-seq correction using mean squared errors, and found that in most cases, the inter-replicate variability was smaller after the correction, or was similar regardless of the correction (**Fig S6**). We then repeated the PCA using ComBat-seq corrected counts and found that samples no longer clustered in a random fashion (**Fig. S3B**). We observed that early time points of all treatments clustered together, and that SP+ samples diverged from V and SP- samples at later time points.

When inspecting each replicate separately using PCA, we observed three outlier samples in replicate C (C_V_6, C_SP+_2 and C_SP+_3; **Fig. S7A-B**). We removed these three samples from the dataset for our final analysis. Time series gene expression data are typically organized along a “U” shape in PCA plots that show the first two principal components (e.g Law et al. 2014). A U-shaped pattern is not seen when we plot all 87 samples, but becomes apparent when plotting one treatment at a time. The U shape is most obvious in virgin females, perhaps because gene expression in virgin females is only influenced by circadian effects, instead of a mix of circadian and mating effects (**Fig. S8A-C**).

### Identification of differentially expressed genes & log_2_ fold change calculation

To identify genes that were significantly differentially expressed between treatments across time, we used three methods (DESeq2 (Love et al. 2014), Limma-Voom (Law et al. 2014) and ImpulseDE2 (Fischer et al. 2018)). These methods differ in how they model variability in the RNA-seq counts, or how they identify dynamic patterns, and we found small differences in their selection of DE genes. DESeq2 models the variability in the RNA-seq counts using a negative binomial distribution, while the Limma-Voom framework models variability in the counts using a Gaussian distribution, after first transforming the counts using precision weights (Law et al. 2014). For both DESeq2 and Limma-Voom, we fit splines with 3 degrees of freedom on the gene expression data. ImpulseDE2 makes use of DESeq2’s framework to model variability in the data, but fits impulse and sigmoid models instead of splines. Once a spline, impulse or sigmoid model is fit for a gene, these methods test whether the estimated model parameters differ across treatments. Using these three methods, we compared RNA abundance between SP+ vs. V, SP- vs. V and SP- vs. SP+. For each of these contrasts, we selected genes as significantly differentially expressed if the gene had *q-*value < 0.05 (Benjamini-Hochberg correction; Benjamini and Hochberg, 1995) in at least two of the three methods.

Next, we also used a log_2_ fold change cutoff to subset these differentially expressed genes. We set up pairwise contrasts in DESeq2 to calculate log_2_ fold changes at each time point, comparing each of the three treatments. We used the *apeglm* method to shrink log_2_ fold change estimates from genes with low overall read counts or highly variable read counts across replicates (Zhu et al. 2019). Genes were selected for our analysis if they had a minimal absolute shrunken fold change of 1.5 or 1.3, at least at one time point (**Table S1**). While shrunken log_2_ fold changes were used to select differentially expressed genes, the original log_2_ fold changes were used in all other analyses and figures.

### Differential exon use analysis

We used DEXSeq (Anders et al. 2012) to quantify differential exon use. First, we aligned fastq files using STAR (Dobin et al. 2013) to the BDGP6.32 genome (the DEXSeq scripts for exon counting are tailored to the ensembl gtf format). We counted exons using DEXSeq scripts and the BDGP6.32.103 gtf. DEXSeq defines exons based on a flattened gene model, i.e. if an exon is present in two or more isoforms, but is shorter in one of the isoforms, the exon is “cut” into two separate exons for the analysis. Next, we filtered out lowly expressed exons and applied batch correction. As a first filtering step, we only kept exons from genes that were analyzed in the gene-level analysis and removed exons that could not be unambiguously assigned to only one gene. This reduced the number of exons from 91,915 to 57,322 exons across 8,375 genes. For the second filtering step, we first normalized exon counts as recommended in Soneson et al. (2016). This normalization step includes normalizing for exon length, gene length, the number of isoforms the gene has and the number of isoforms the exon is part of. Then, we removed 20% of exons with the lowest normalized counts. This further reduced the number of exons to 48,822 exons across 7,671 genes. At this point, we corrected the filtered raw exon counts for the immune response batch effect using Combat-seq (Zhang et al. 2020), as done for the gene-level analysis. We found that the most significant correction of exon counts occurred in the exons of genes that encode antimicrobial peptides. We also compared inter-replicate variability before and after Combat-seq correction using mean squared errors, and found that in most cases, the inter-replicate variability was smaller after the correction (**Fig S9**). After batch correction, as a final filtering step, we removed exons that are shared across all transcript isoforms of their gene, or that are present in a gene that has only one annotated transcript isoform. This gave us a final set of 16,011 exons across 4,143 genes for the differential exon use analysis. As for the gene-level analysis, we removed three outlier samples (C_V_6, C_SP+_2 and C_SP+_3).

The DEXSeq R package (Anders et al. 2012) assesses differential exon use between treatments while taking into account gene-level differential expression. We set up one DEXSeq test per time point per exon, each time comparing two treatments. We selected exons with significant differential use based on 1) a *q*-value < 0.05 (Benjamini-Hochberg correction); 2) a gene level *q-value* < 0.05 (calculated using the DEXSeq function perGeneQValue) and 3) a minimum 50% fold difference in exon use.

When we inspected differential exon use by plotting variance stabilized normalized exon counts in function of time, some of the selected exons appeared to be potential false positives. Suspected false positives demonstrated a high inter-replicate variability at many time points, except at the time point at which significant differential use was detected. To differentiate between likely and unlikely false positives, we made use of the time series data: For each exon and for each treatment separately, we fit a natural cubic spline function through all the data points (normalized counts across 3 replicates over time). We then selected only exons whose spline fit had R^2^ > 0.6 for at least one of the three treatments (a good spline fit suggests a higher consistency among the replicates across time). The final selection of differentially expressed exons is listed in **Table S2**. This table also contains the start and end location of each exon.

### Cluster analysis

For all pairs of differentially expressed features (genes and exons), we determined the similarity of their expression profiles, represented as *log_2_(SP+/V)*, using the LLR^2^ method (Venkatraman et al. 2021). This method uses empirical Bayesian regression and prior biological evidence of association to fit ordinary differential equation models to temporal feature profiles. These pairwise similarity scores are used to perform hierarchical clustering on the set of temporal gene trajectories. The purpose of using differential equation-based similarity measures for clustering is to more reliably capture similarities in the nonlinear temporal dynamics of two genes, which are explicitly modeled by such equations, compared to more common similarity measures such as Pearson correlation (Farina et al. 2008). Pairwise similarity measures between genes are derived from these models, and are shrunken if there is no prior evidence that both genes interact.

We obtained prior biological evidence from three sources. First, we used gene-gene interaction scores from the STRING database (v11.0) (Szklarczyk et al. 2019). Gene pairs with STRING scores ≥ 900 (assigned as “very high confidence” in the STRING database) were given prior = 1. Gene pairs with STRING scores < 900 and ≥ 700 (“high confidence”) were given prior = NA. Second, we assigned priors based on correlations in replicate datasets. Within each of our 3 replicate datasets, we calculated correlations on the normalized counts using LPWC (default settings without lag; (Chandereng and Gitter 2020). If a feature pair had an average correlation across the 3 replicate datasets ≥ 0.9, the pair received prior = 1. If the average correlation was < 0.9 and ≥ 0.8, the pair received prior = NA. Third, we assigned prior = NA to features that were predicted to be expressed in the same cell type based on deconvolution analysis (see below). The prior of any feature pair that received prior = NA from more than one prior source was set to 1. All other pairs received prior = 0. Of all feature pairs in our dataset, 94% had prior = 0, 4.59% had prior = NA and 1.41% had prior = 1. We clustered features using hierarchical clustering (method Ward.D), using 1-LLR^2^ as a representation of the distance between features. In time series datasets, often many genes have highly similar expression profiles. The goal of prior incorporation is to put more emphasis on genes that are likely functionally related, e.g. by functioning in the same biological pathway, or by being expressed in the same cell type. At the same time, priors based on replicate correlations still allow for novel, data-driven discoveries, as illustrated in **figure S10**.

For the cluster analysis, we removed 3 exons (*Eip75B:E001, sNPF-R:E004, Sox120F:E006*) and 2 genes (*Dro, Amy-p*) with irregular expression patterns, and 5 additional genes (*Paics, CG31370, CG31664, apolpp, asRNA:CR45175*) which did not have a single *q-value* < 0.05 at any time point, for any treatment comparison, according to pairwise comparisons in DESeq2 (even though they were identified as differentially expressed using the temporal differential expression methods we applied). Gene and exon cluster assignment can be found in **Table S4**.

### Deconvolution of bulk RNA-seq using single cell RNA-sequencing data

To determine in which cell types differential expression might occur, we deconvoluted our bulk RNA-seq dataset using single cell RNA-seq of the head as a reference. We downloaded single cell RNA-seq of the *Drosophila* head from Scope (10x, stringent; Li et al. 2021). From this dataset, we selected only cells from females and used the available annotations (“annotation_broad”) that assign cells to annotated cell types. Cells annotated with “artefact” were removed. We processed this dataset using Seurat V4 (Hao et al. 2021). Briefly, we only kept features expressed in at least 3 cells, and only kept cells that expressed at least 200 features. We normalized and scaled the data and used the FindMarkers function to compare each of the annotated cell types with all other cell types in the dataset. The resulting marker genes, log_2_ fold changes and p-values were used to create a signature matrix for the R package scMappR (Sokolowski et al. 2021). scMappR uses this signature matrix to estimate cell type proportions in bulk RNA-seq data. In addition, scMappR calculates cell-weighted fold changes for differentially expressed genes. These can be used to estimate in which of the cell types the differential expression is most likely to occur. We ran scMappR using default settings on each time point, each time comparing mated (SP+) and virgin females. We found that cell type proportions did not differ significantly between mated (SP+) and virgin females at any time point. For a subset of the differentially expressed genes, scMappR was able to calculate a cell type specificity score, indicating how likely it is that the gene is undergoing expression changes in a specific cell type. We assigned these genes to specific cell types based on their maximal specificity value at any time point, requiring a minimum absolute specificity of 0.5. This allowed us to assign 386 differentially expressed genes to an annotated cell type (**Table S5)**.

### Functional interpretation of results and motif enrichment analysis

To investigate biological functions of differentially expressed genes, we ran GO enrichment analyses using ClusterProfiler and rrvgo (Ashburner et al. 2000; The Gene Ontology Consortium 2021; Wu et al. 2021; Yu et al. 2012; Sayols et al. 2021). All GO terms listed in this study are significantly enriched with a *q-*value < 0.05 (Benjamini-Hochberg correction) and based on a minimum gene set size of 10. We consulted Flybase (FB2021_01; Larkin et al. 2021) to acquire additional information for differentially expressed genes of interest. FlyAtlas 2 (Leader et al. 2018) was used to obtain tissue-specific gene expression measurements. RcisTarget was used for transcription factor binding motif enrichment analysis (Aibar et al. 2017). We applied default settings and used all 9,027 expressed genes in our dataset as background. We used the database “dm6-5kb-upstream-full-tx-11species.mc8nr” to search for motifs in a 5 kb region upstream of transcription start sites. RcisTarget was run on each cluster defined by the LLR^2^ method. Of all transcription factors that were identified using RcisTarget (both using direct inference and based on homology), we only kept ones that were differentially expressed in our dataset, had at least 3 predicted targets, and whose targets had a median absolute LPWC (Chandereng and Gitter 2020) correlation ≥ 0.8. From the resulting set of transcription factors, we manually selected ones that are up- or downregulated around the same time or earlier than their predicted targets.

### Identification of genes with circadian rhythms

We identified genes with circadian rhythms in virgin females using the R package Metacycle (Wu et al. 2016). Minimum periodicity was set to 22 hours; maximum periodicity was set to 26 hours. We extracted results from the LS algorithm and classified genes with a *q-value* < 0.3 (Benjamini-Hochberg correction) as having a circadian rhythm in virgin females.

### R packages used to generate figures

Figures were made using base R (R version 4.1.0) and the R packages ComplexHeatmap (Gu et al. 2016), igraph (Scardi et al. 2006), eulerr (Wilkinson 2012) and ggplot2 (Wilkinson 2012; Wickham 2009).

### Querying of the fly hemibrain connectome

We investigated connections between pC1 (a, b and c) neurons and neurons with roles in circadian behaviors using the “shortest path” tool in the connectome neuprint (hemibrain v1.2.1; Scheffer et al. 2020), using the default minimum of 10 synapses. We recorded the number of interneurons for each path as 0 (direct connection), 1 (1 interneuron), 2 (2 interneurons) or >2 (more than 2 interneurons) (**Table S6**). For paths with 2 or fewer interneurons, we registered whether SMP454 or oviIN neurons were used as interneurons.

## Supplemental Figures

**Figure S1:**
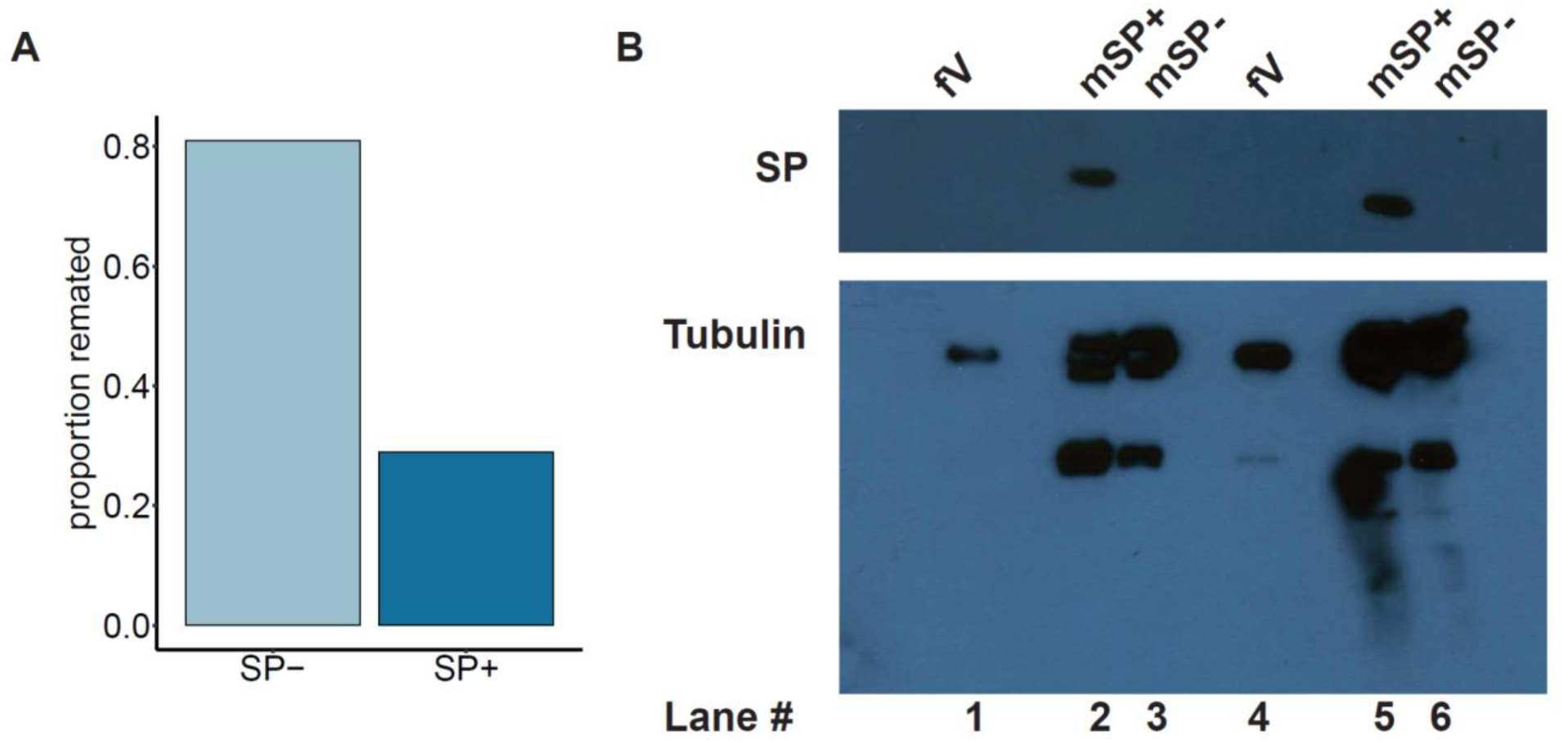
Verification of the absence of SP in SP null males. A) Proportion of remated females in a four day receptivity assay, after mating to a *SP* null male (SP-) or a control male (SP+). Fisher’s exact test *p-value* = 0.009. *n* re-mated for SP- = 13/16 females; *n* re-mated for SP+ = 4/14 females. B) Western blot showing the presence of SP in control males (mSP+; *n* = 1 accessory gland) and the absence of SP in SP null males (mSP-; *n* = 1 accessory gland). Virgin female reproductive tracts (fV; *n* = 4 tracts) act as a negative control. Tubulin serves as a loading control.

**Figure S2:**
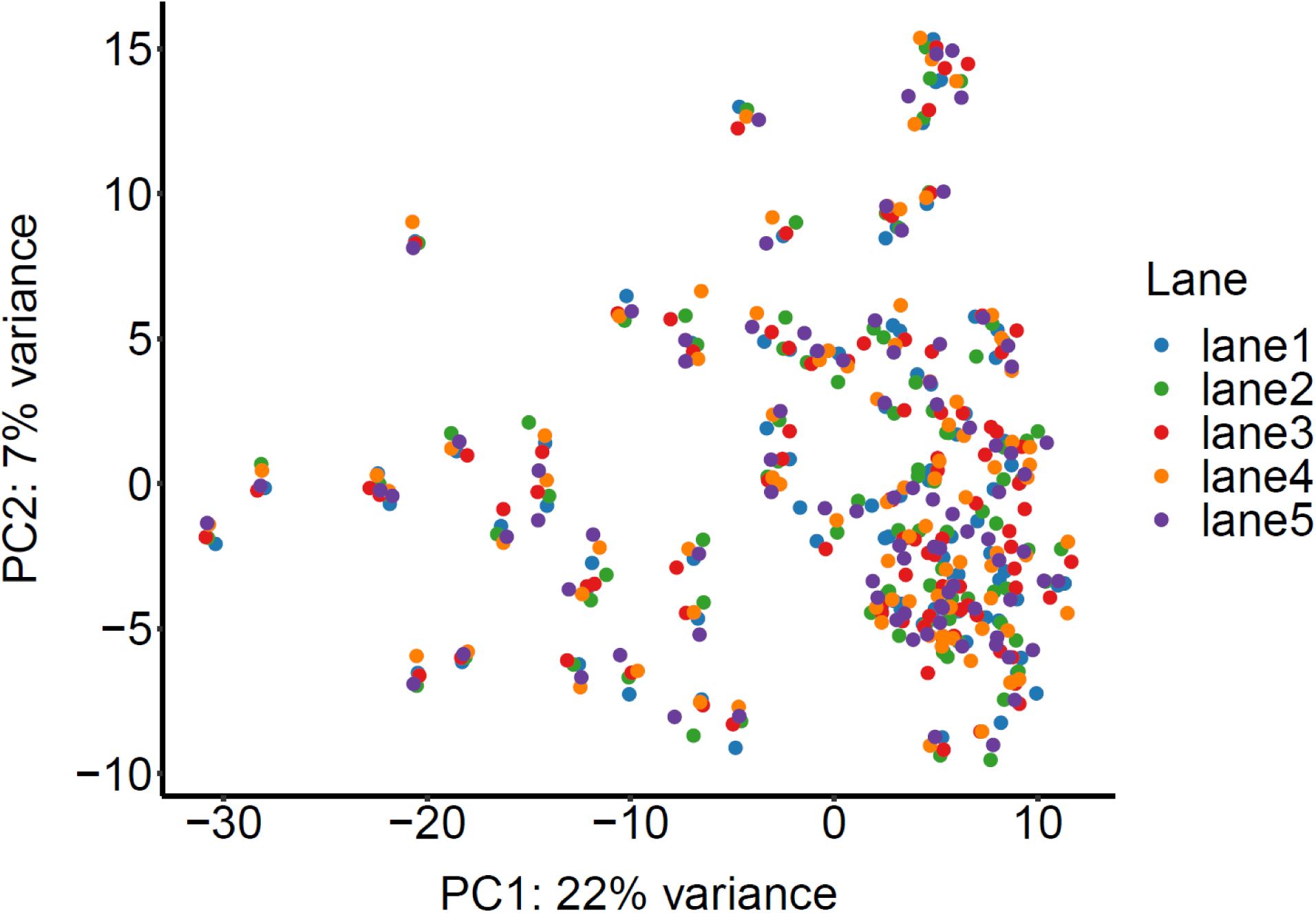
Principal component plot based on analysis using counts from 450 fastq files (90 samples sequenced on 5 lanes). PCA was used to investigate potential lane effects.

**Figure S3:**
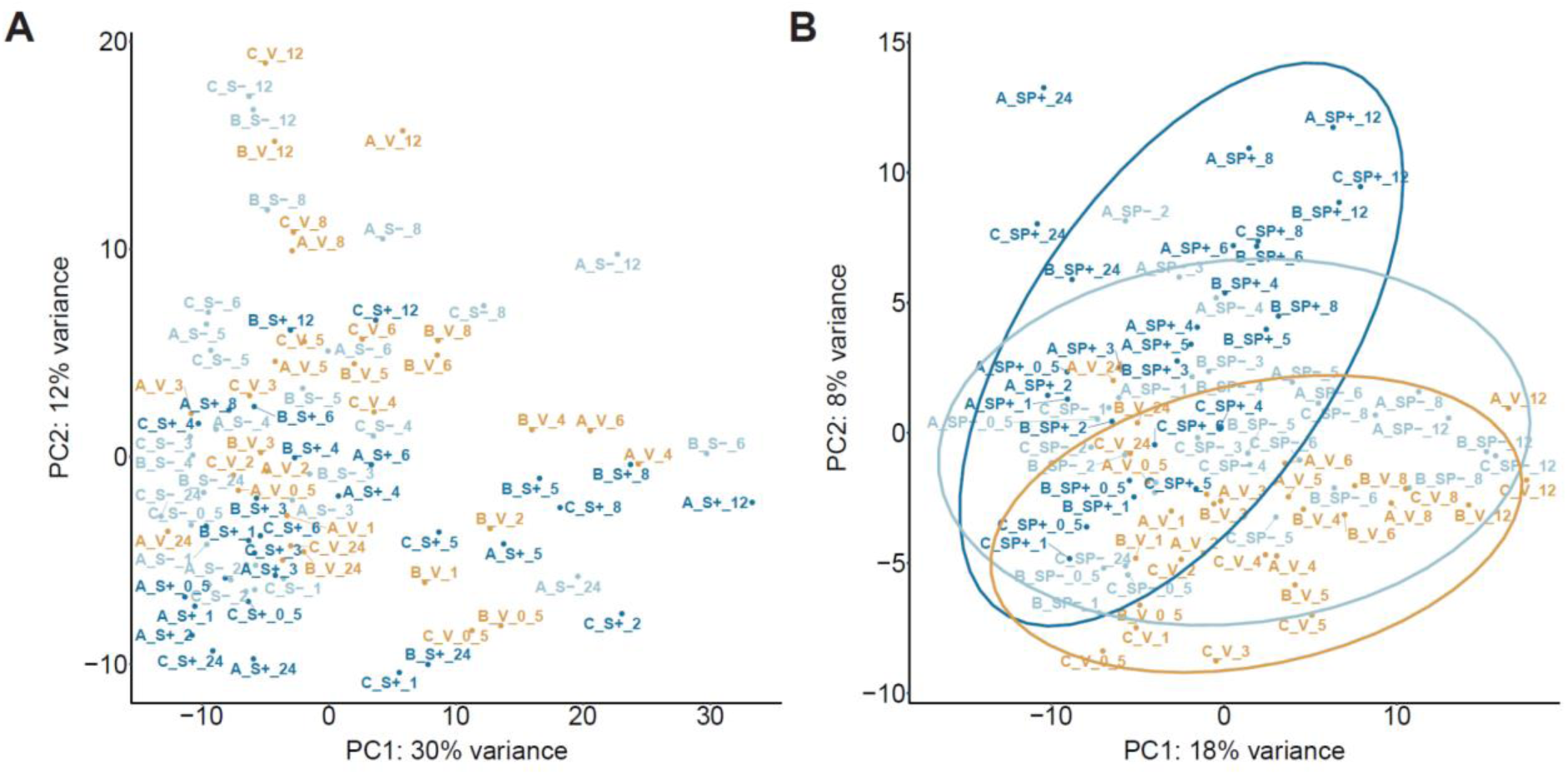
Principal component plot based on the 500 most variable genes, before and after ComBat-seq correction. A) Principal component plot based on analysis across all samples before ComBat-seq batch correction. B) Principal component plot based on analysis across all samples after ComBat-seq batch correction. Sample notation: replicate_treatment_timepoint

**Figure S4:**
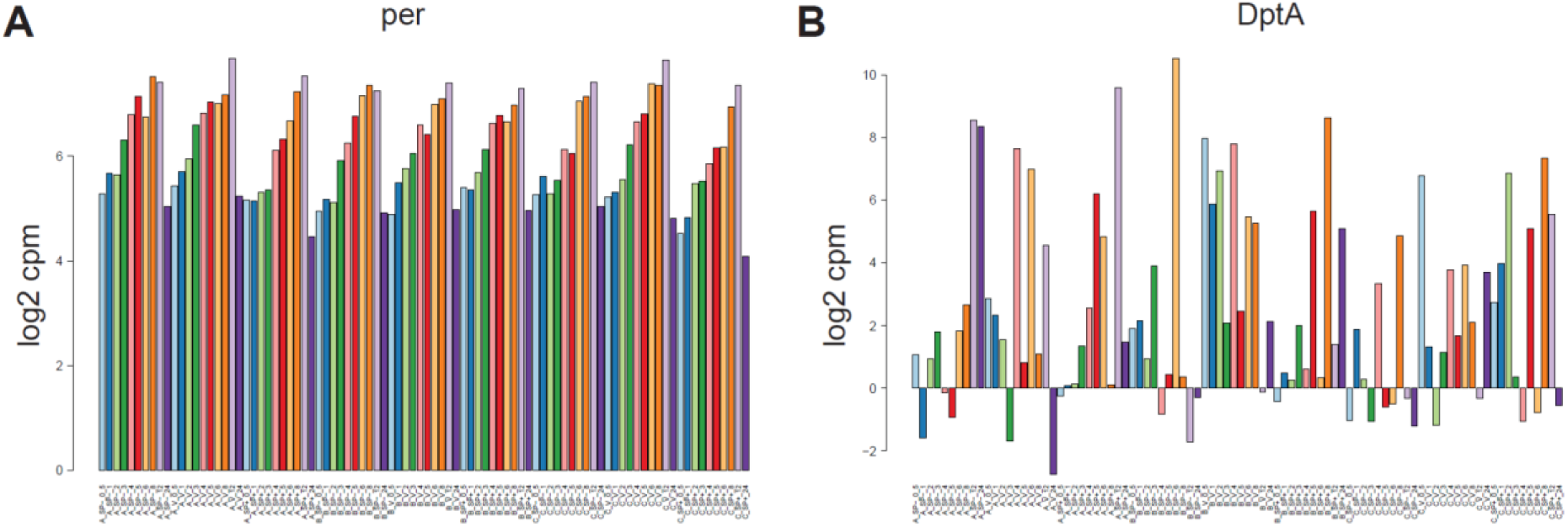
Examples of genes that are among the top 100 most variable genes in the dataset. A) Example of a gene with a circadian rhythm, with regular expression patterns across samples. B) Example of an immune response gene, with irregular expression patterns across samples.

**Figure S5:**
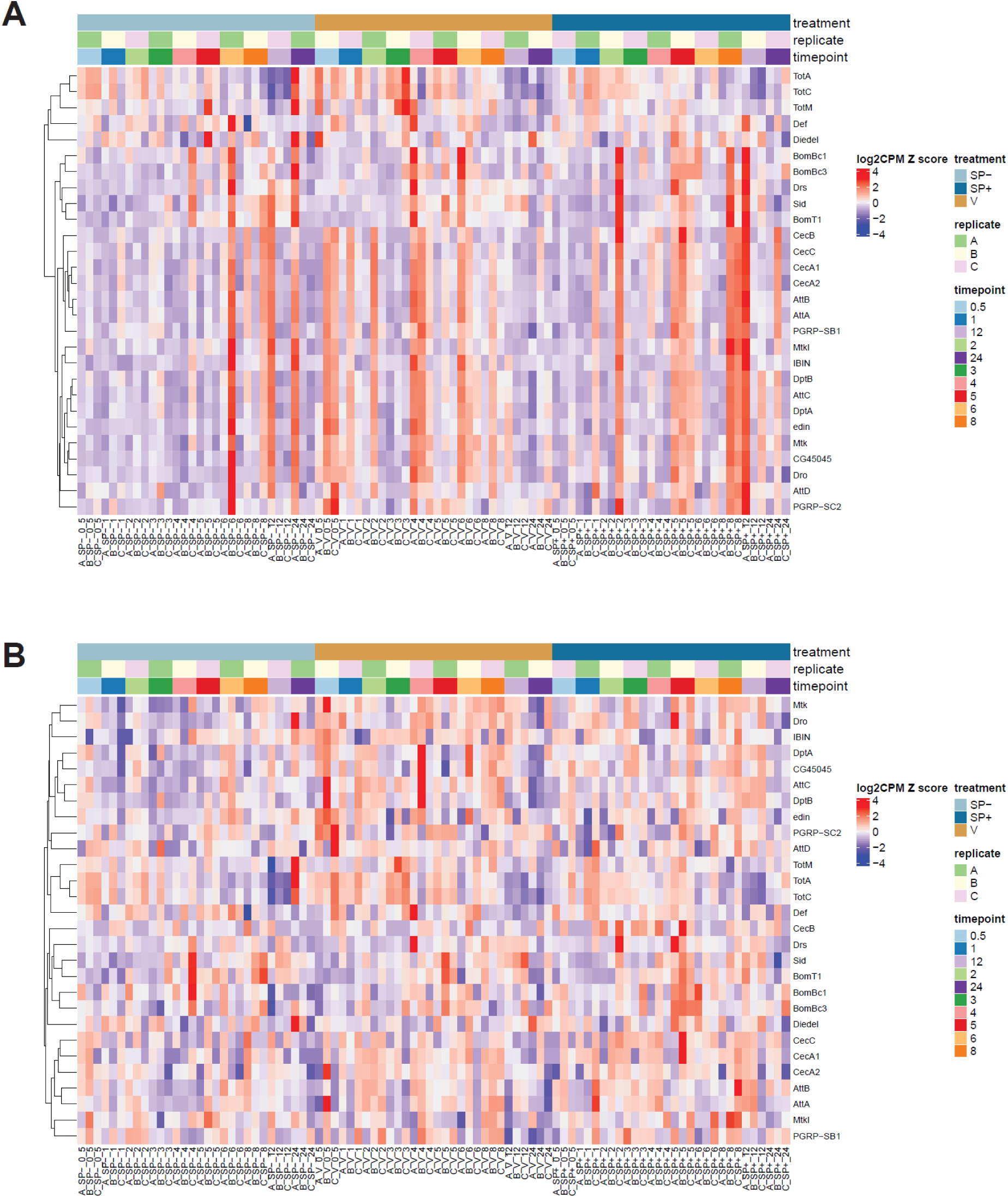
Heatmaps showing original and ComBat-seq batch-corrected counts for 28 immune genes. A) Heatmap showing the Z scores of log_2_-transformed counts per million (CPM) for 28 immune genes across all samples, before batch correction. B) Heatmap showing the Z scores of log_2_-transformed counts per million (CPM) for 28 immune genes across all samples after ComBat-seq correction. Dendrograms show hierarchical clustering of genes (rows) based on Euclidean distance.

**Figure S6:**
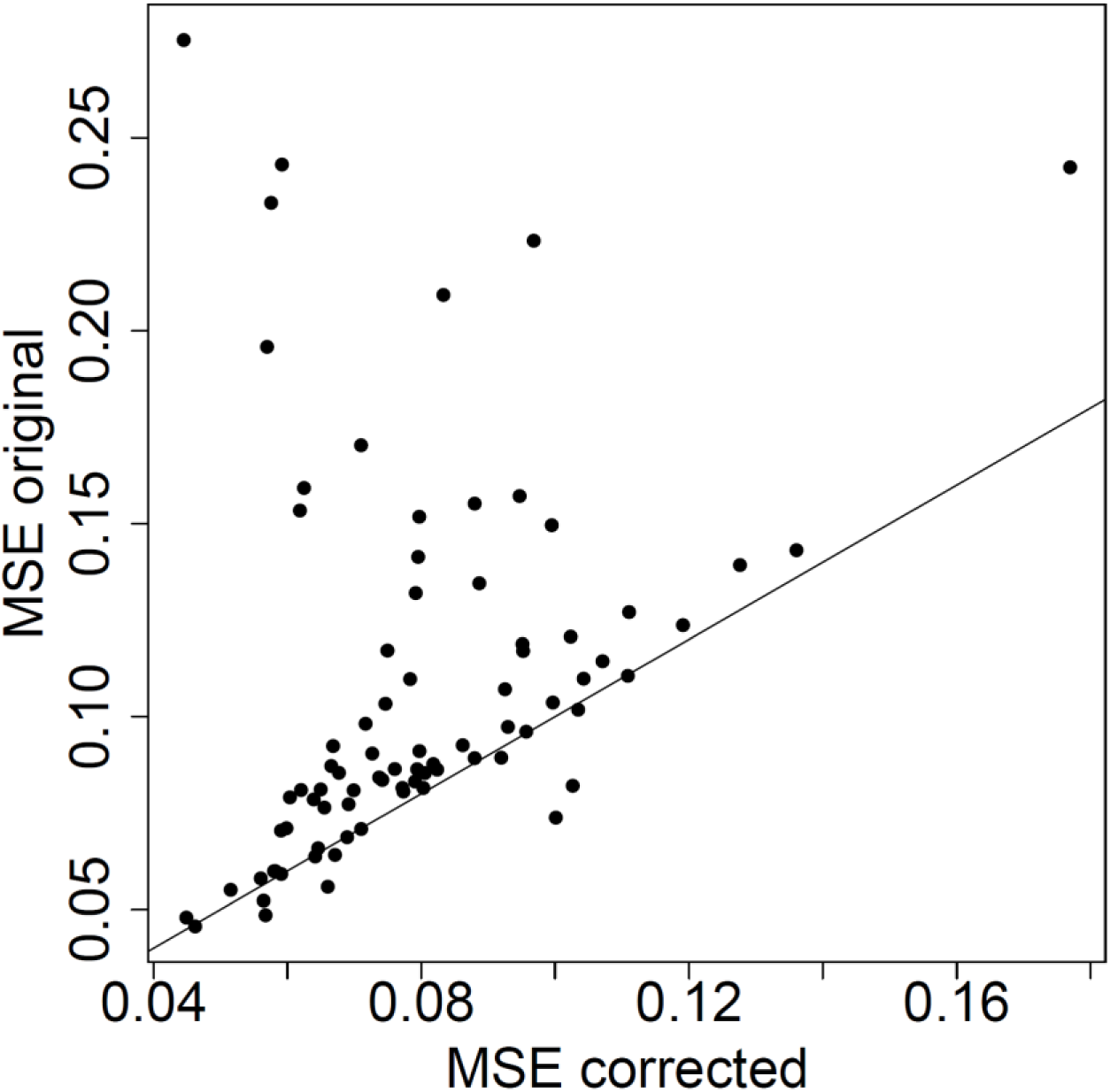
Mean Squared Error (MSE) between replicates, calculated using original and batch-corrected log_2_-transformed normalized counts per million. Each dot represents the comparison between two replicates.

**Figure S7:**
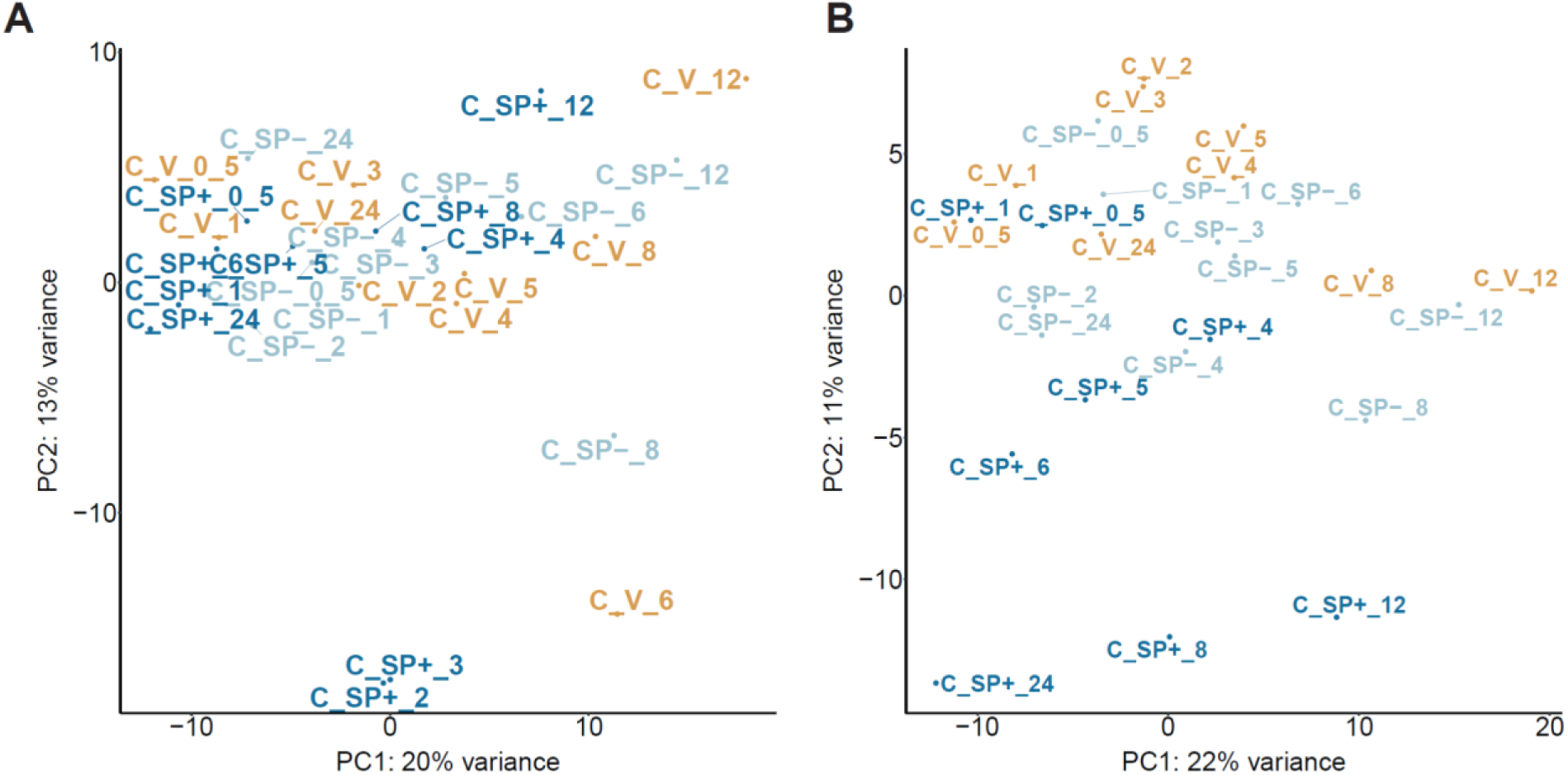
Principal component plots based on analysis across all samples in replicate C, using the top 500 genes with highest row variance after ComBat-seq batch correction, before and after removal of outlier samples. A) Three samples are outliers relative to other samples in replicate C (C_SP+_3, C_SP+_2, C_V_6). B) PC plot after removal of three outlier samples. Sample notation: replicate_treatment_timepoint

**Figure S8:**
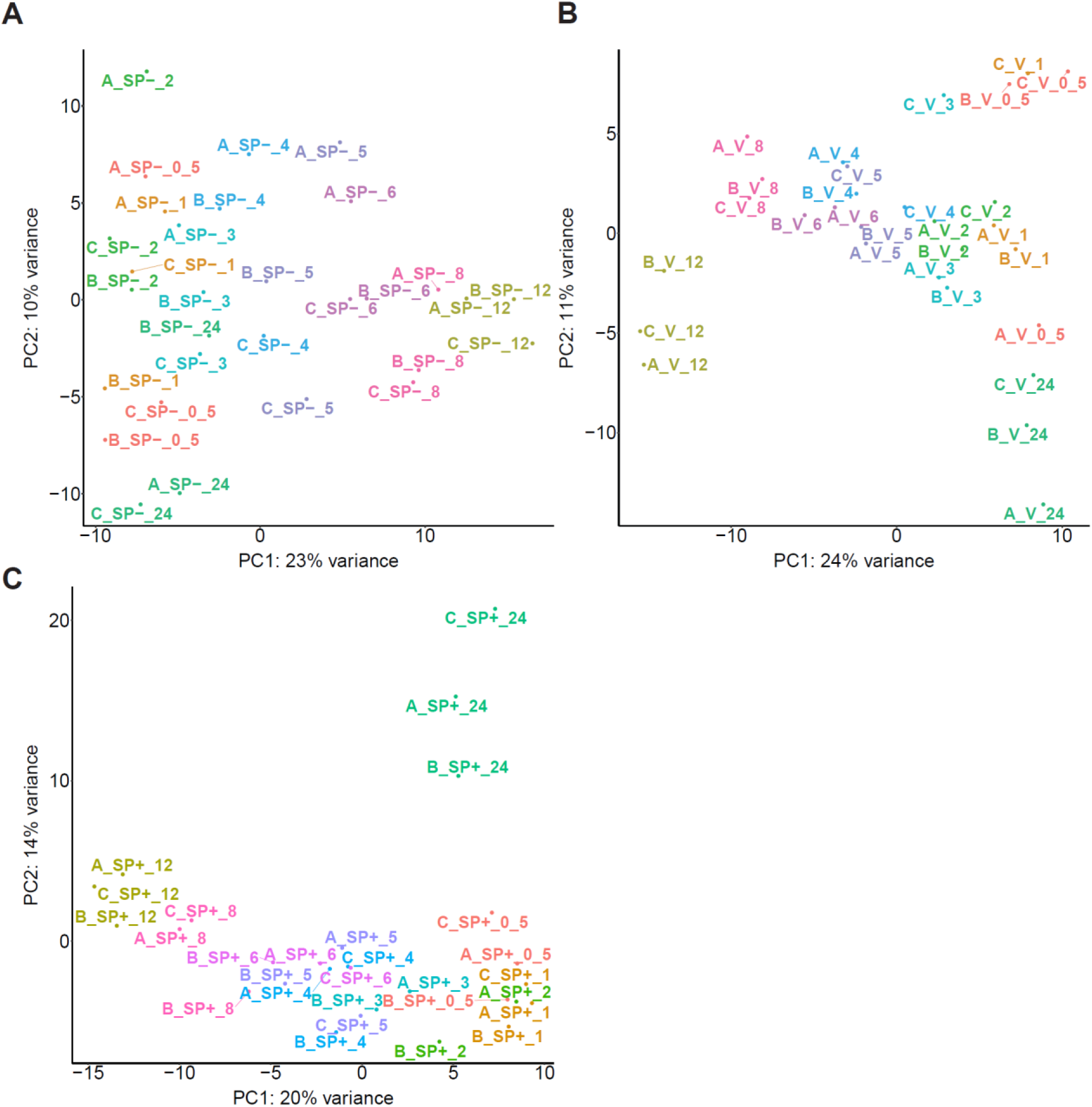
Principal component plots based on analysis across all samples per treatment, using the top 500 genes with highest row variance, after ComBat-seq batch correction. A) PC plot for females mated to *SP* null males (SP-). B) PC plot for virgin females (V). C) PC plot for females mated to control males (SP+). Sample notation: replicate_treatment_timepoint

**Figure S9:**
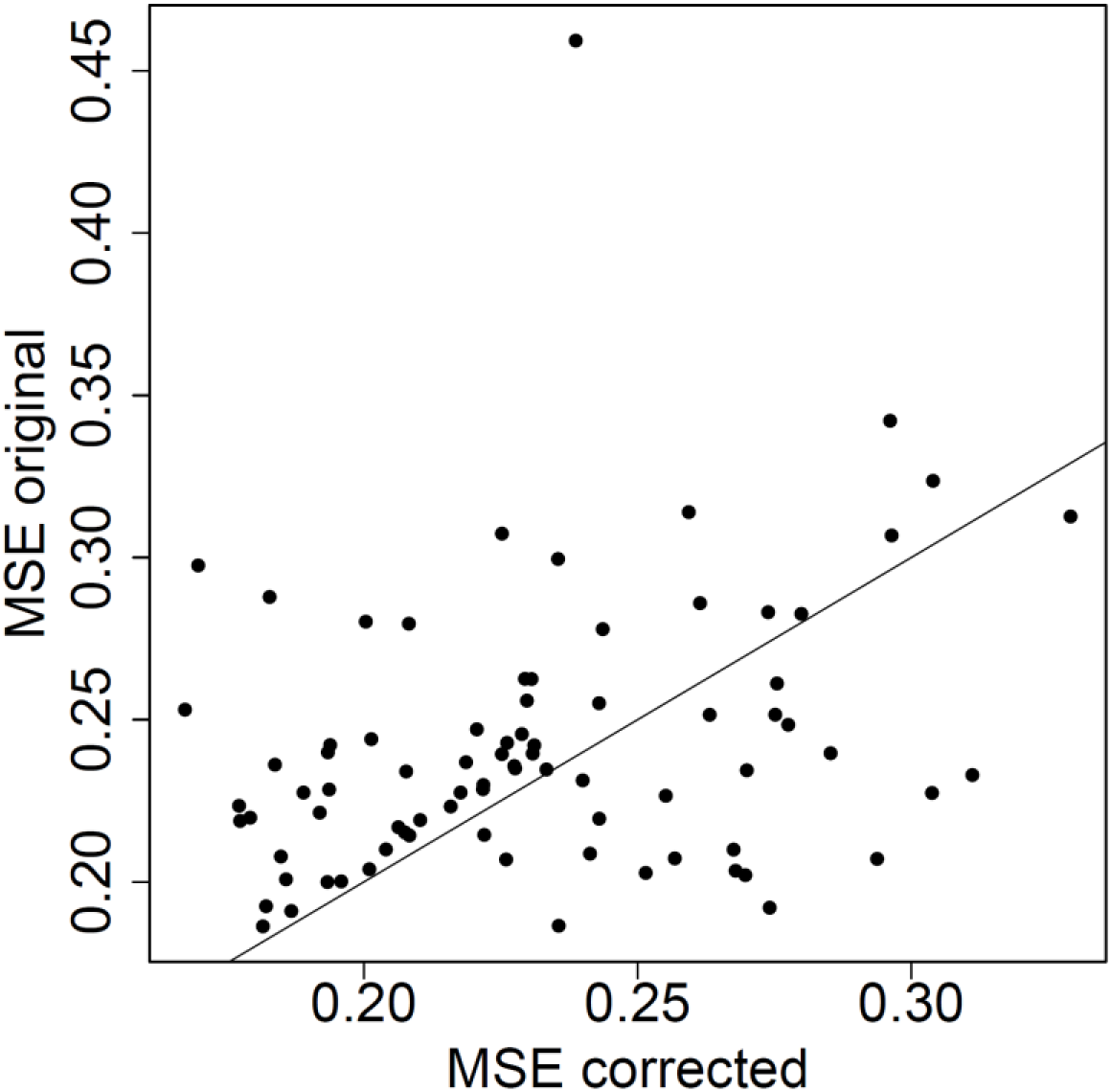
Mean Squared Error (MSE) between replicates, calculated using original and batch-corrected exon-level log_2_-transformed normalized counts. Each dot represents the comparison between two replicates.

**Figure S10:**
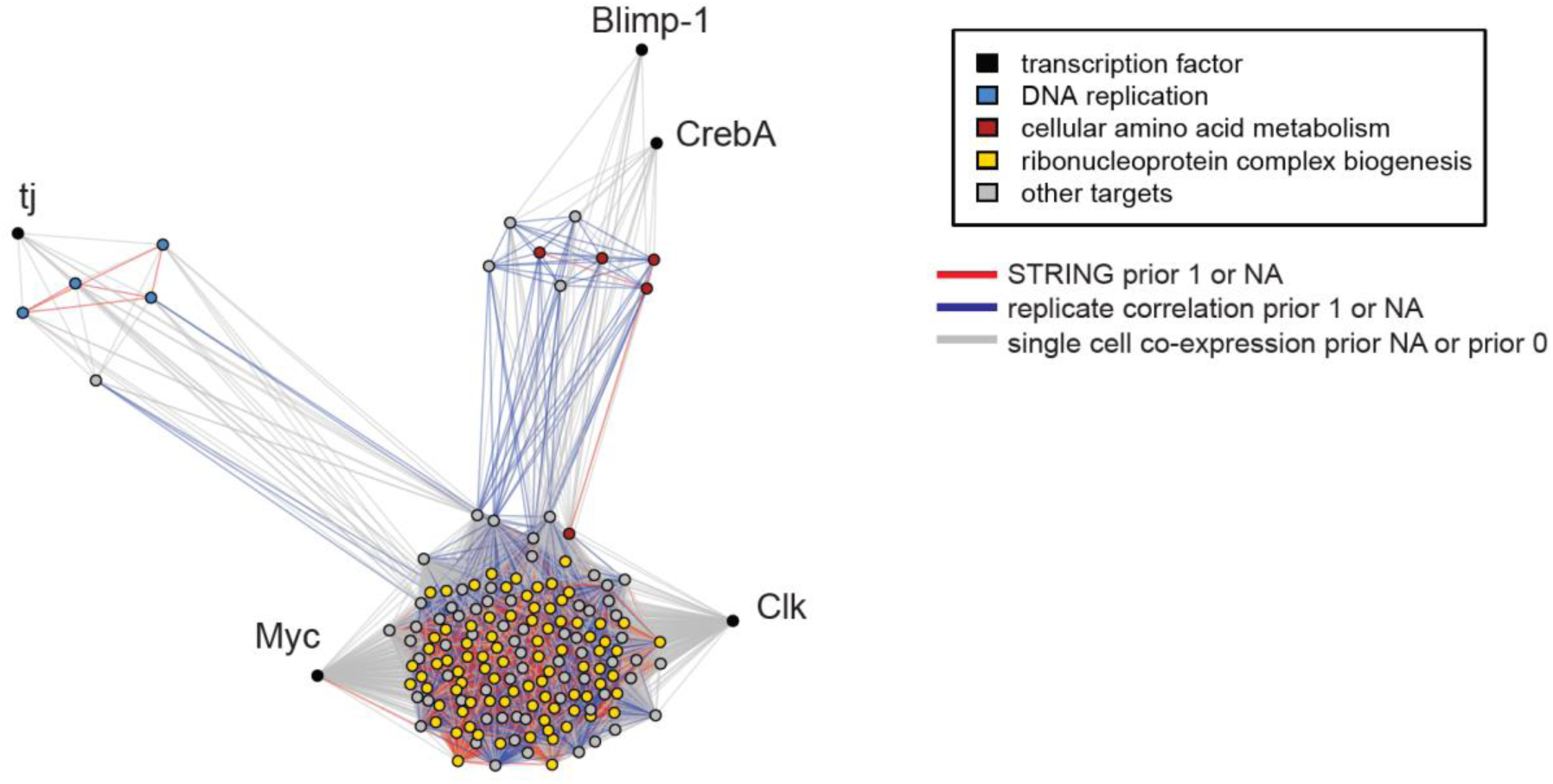
Network showing transcription factors and predicted target genes. Edges represent distances between features based on LLR^2^ values. Only edges are shown between transcription factors and predicted targets, and among predicted targets of the same transcription factor. Nodes are colored based on significant GO term enrichment. Edges are colored based on prior value. This figure illustrates that close clustering of genes is not driven solely by STRING based prior knowledge.

**Figure S11:**
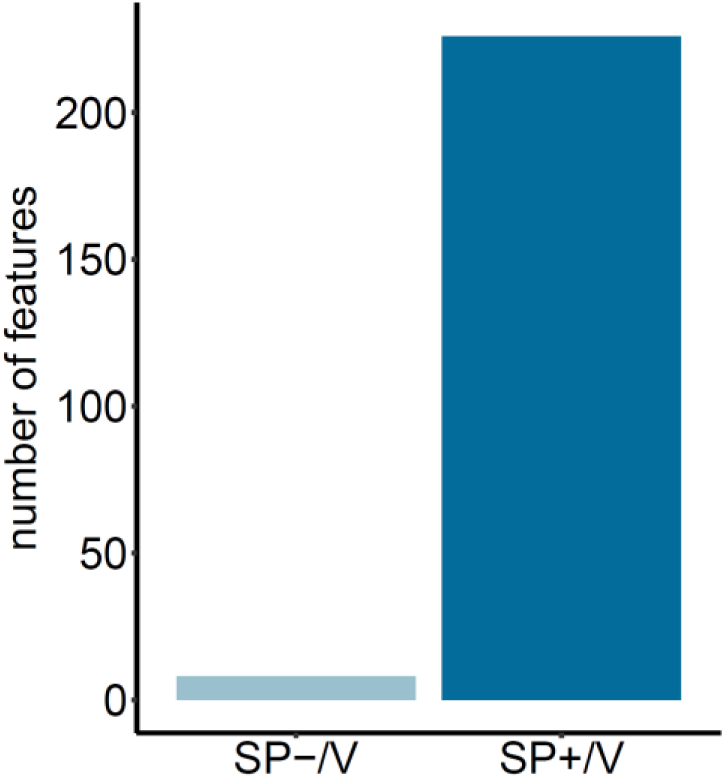
Number of genes and exons that were significantly differentially expressed within 30 minutes to 4 hours post-mating, and that are still differentially expressed at any time between 5 hours and 24 hours post-mating. Significant differential expression is based on a *q-*value < 0.05, in mated relative to virgin (V) females in the absence (SP-) or presence (SP+) of SP.

**Figure S12:**
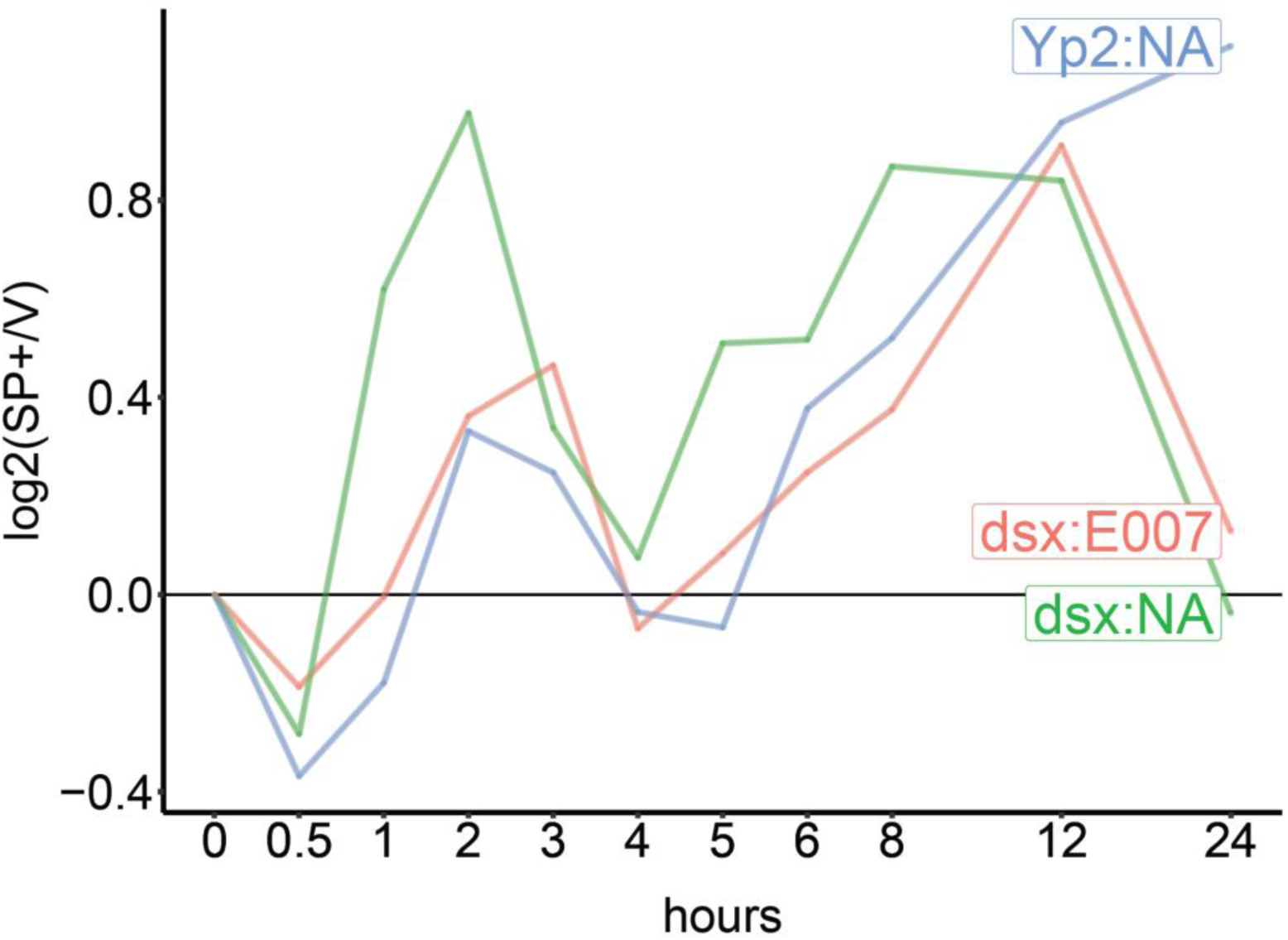
Time series log_2_ fold changes of mated (SP+) versus virgin (V) females for *dsx* gene and *dsx* exon *007,* and *Yolk protein 2* (*Yp2*).

**Figure S13:**
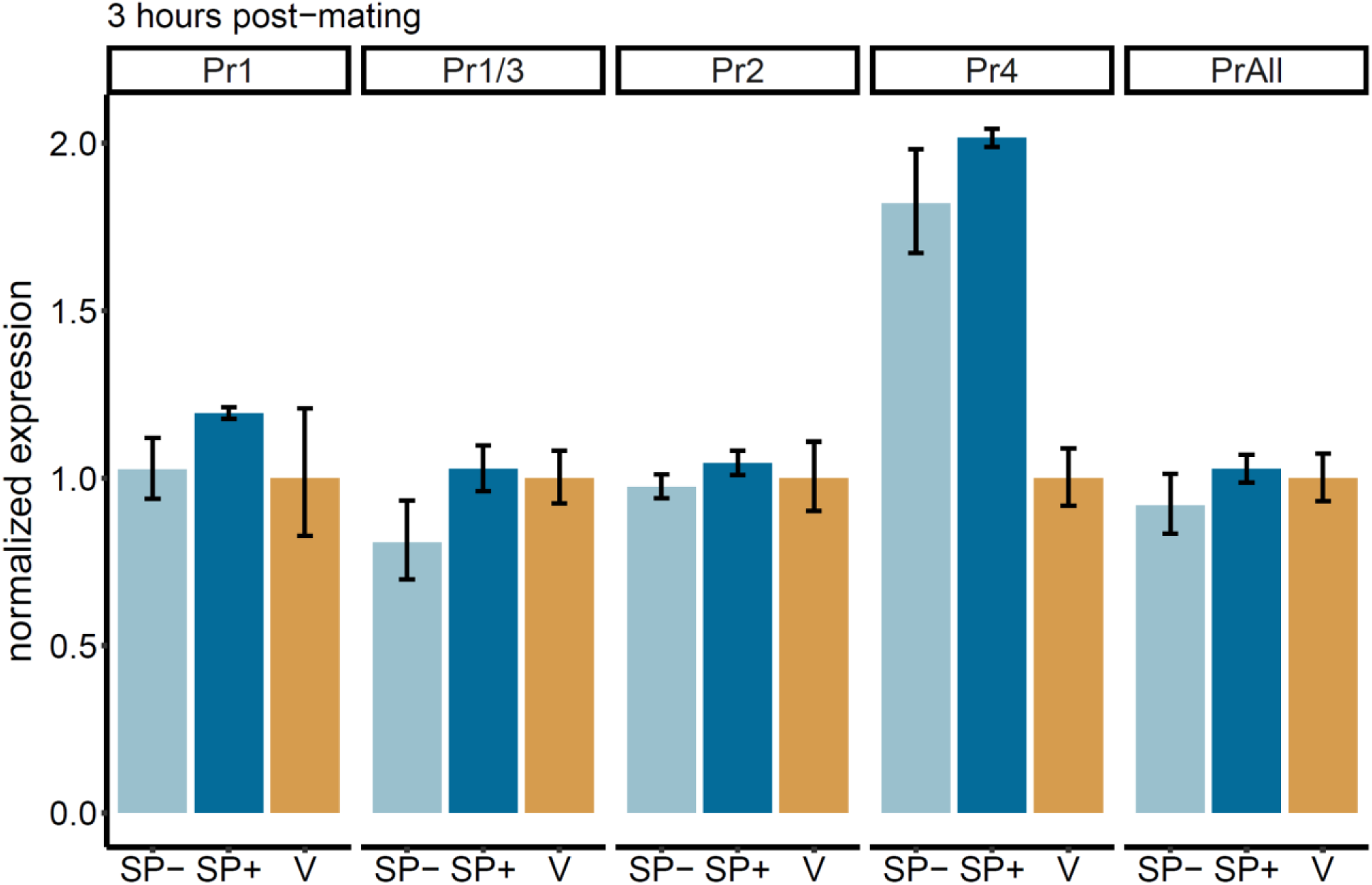
Post-mating expression of transcripts controlled by 4 promoters of the *foraging* gene, measured by qPCR. Error bars represent standard error of the mean based on 3 biological replicates. Pr1 = promoter 1; Pr2 = promoter 2; Pr4 = promoter 4; Pr1/3 captures expression of transcripts controlled by promoters 1 and 3, since transcripts from promoter 3 are nested within those of promoter 1. PrAll captures the expression of exons that are part of all *foraging* transcripts.

## References

Abruzzi, Katharine Compton, Joseph Rodriguez, Jerome S. Menet, Jennifer Desrochers, Abigail Zadina, Weifei Luo, Sasha Tkachev, and Michael Rosbash. 2011. “*Drosophila* CLOCK Target Gene Characterization: Implications for Circadian Tissue-Specific Gene Expression.” Genes & Development 25 (22): 2374–86.

Aibar, Sara, Carmen Bravo González-Blas, Thomas Moerman, Vân Anh Huynh-Thu, Hana Imrichova, Gert Hulselmans, Florian Rambow, et al. 2017. “SCENIC: Single-Cell Regulatory Network Inference and Clustering.” Nature Methods 14 (11): 1083–86.

Akagi, Kazutaka, Moustafa Sarhan, Abdel-Rahman S. Sultan, Haruka Nishida, Azusa Koie, Takumi Nakayama, and Hitoshi Ueda. 2016. “A Biological Timer in the Fat Body Comprising Blimp-1, βFtz-f1 and Shade Regulates Pupation Timing in *Drosophila Melanogaster*.” Development 143 (13): 2410–16.

Allemand, R. 1976. “Influence of Light Condition Modification on the Circadian Rhythm of Vitellogenesis and Ovulation in *Drosophila Melanogaster*.” Journal of Insect Physiology 22 (8): 1075–80.

Allen, Aaron M., Ina Anreiter, Megan C. Neville, and Marla B. Sokolowski. 2017. “Feeding-Related Traits Are Affected by Dosage of the Foraging Gene in *Drosophila Melanogaster*.” Genetics. https://doi.org/10.1534/genetics.116.197939.

Anders, Simon, Paul Theodor Pyl, and Wolfgang Huber. 2015. “HTSeq--a Python Framework to Work with High-Throughput Sequencing Data.” Bioinformatics 31 (2): 166–69.

Anders, Simon, Alejandro Reyes, and Wolfgang Huber. 2012. “Detecting Differential Usage of Exons from RNA-Seq Data.” Genome Research 22 (10): 2008–17.

Anreiter, Ina, Jamie M. Kramer, and Marla B. Sokolowski. 2017. “Epigenetic Mechanisms Modulate Differences in *Drosophila* Foraging Behavior.” Proceedings of the National Academy of Sciences. https://doi.org/10.1073/pnas.1710770114.

An, W., and P. C. Wensink. 1995. “Integrating Sex- and Tissue-Specific Regulation within a Single *Drosophila* Enhancer.” Genes & Development 9 (2): 256–66.

Apger-McGlaughon, Jennifer, and Mariana F. Wolfner. 2013. “Post-Mating Change in Excretion by Mated *Drosophila Melanogaster* Females Is a Long-Term Response That Depends on Sex Peptide and Sperm.” Journal of Insect Physiology 59 (10): 1024–30.

Ashburner, Michael, Catherine A. Ball, Judith A. Blake, David Botstein, Heather Butler, J. Michael Cherry, Allan P. Davis, Kara Dolinski, Selina S. Dwight, Janan T. Eppig, Midori A. Harris, David P. Hill, Laurie Issel-Tarver, Andrew Kasarskis, Suzanna Lewis, John C. Matese, Joel E. Richardson, Martin Ringwald, Gerald M. Rubin, Gavin Sherlock. 2000. “Gene Ontology: Tool for the Unification of Biology. The Gene Ontology Consortium.” Nature Genetics 25(1):25–29 doi: 10.1038/75556

Avila, Frank W., Alexandra L. Mattei, and Mariana F. Wolfner. 2015. “Sex Peptide Receptor Is Required for the Release of Stored Sperm by Mated *Drosophila Melanogaster* Females.” Journal of Insect Physiology 76 (May): 1–6.

Avila, Frank W., K. Ravi Ram, Margaret C. Bloch Qazi, and Mariana F. Wolfner. 2010. “Sex Peptide Is Required for the Efficient Release of Stored Sperm in Mated *Drosophila Females*.” Genetics. https://doi.org/10.1534/genetics.110.119735.

Bastock, Margaret and Aubrey Manning. 1955. “The courtship of *Drosophila melanogaster*.” Behaviour 8, 85–111.

Bath, Eleanor, Samuel Bowden, Carla Peters, Anjali Reddy, Joseph A. Tobias, Evan Easton-Calabria, Nathalie Seddon, Stephen F. Goodwin, Stuart Wigby. 2017. “Sperm and Sex Peptide Stimulate Aggression in Female *Drosophila*.” Nature ecology & evolution, 1(6), 0154. https://doi.org/10.1038/s41559-017-0154

Bejamini Yoav, Yosef Hochberg. 1995. “Controlling the False Discovery Rate: A Practical and Powerful Approach to Multiple Testing.” Journal of the Royal Statistical Society: Series B (methodological) 57 (1): 289–300.

Bolger, Anthony M., Marc Lohse, and Bjoern Usadel. 2014. “Trimmomatic: A Flexible Trimmer for Illumina Sequence Data.” Bioinformatics 30 (15): 2114–20.

Bontonou, Gwénaëlle, Haq Abdul Shaik, Béatrice Denis, and Claude Wicker-Thomas. 2015. “Acp70A Regulates *Drosophila Pheromones* through Juvenile Hormone Induction.” Insect Biochemistry and Molecular Biology 56 (January): 36–49.

Bousquet, François, Tetsuya Nojima, Benjamin Houot, Isabelle Chauvel, Sylvie Chaudy, Stéphane Dupas, Daisuke Yamamoto, and Jean-François Ferveur. 2012. “Expression of a Desaturase Gene, *desat1*, in Neural and Nonneural Tissues Separately Affects Perception and Emission of Sex Pheromones in *Drosophila*.” Proceedings of the National Academy of Sciences of the United States of America 109 (1): 249–54.

Bownes, M., A. Scott, and A. Shirras. 1988. “Dietary Components Modulate Yolk Protein Gene Transcription in *Drosophila Melanogaster*.” Development 103 (1): 119–28.

Bradley, T. J., and F. H. Simmons. 1997. “An Analysis of Resource Allocation in Response to Dietary Yeast in *Drosophila Melanogaster*.” Journal of Insect Physiology 43 (8): 779–88.

Bromfield, John J. 2014. “Seminal Fluid and Reproduction: Much More Than Previously Thought.” Journal of assisted reproduction and genetics, 31(6), 627–636. https://doi.org/10.1007/s10815-014-0243-y

Burtis, K. C., K. T. Coschigano, B. S. Baker, and P. C. Wensink. 1991. “The Doublesex Proteins of *Drosophila Melanogaster* Bind Directly to a Sex-Specific Yolk Protein Gene Enhancer.” The EMBO Journal 10 (9): 2577–82.

Carney, G. E., and M. Bender. 2000. “The *Drosophila Ecdysone Receptor* (*EcR*) Gene Is Required Maternally for Normal Oogenesis.” Genetics 154 (3): 1203–11.

Carvalho, Gil B., Pankaj Kapahi, David J. Anderson, and Seymour Benzer. 2006. “Allocrine Modulation of Feeding Behavior by the Sex Peptide of *Drosophila*.” Current Biology: CB 16 (7): 692–96.

Cavanaugh, Daniel J., Jill D. Geratowski, Julien R.A. Wooltorton, Jennifer M. Spaethling, Clare E. Hector, Xiangzhong Zhen, Erik C. Johnson, James H. Eberwine, Amita Sehgal. 2014. “Identification of a Circadian Output Circuit for Rest:Activity Rhythms in *Drosophila*.” Cell 157 (3): 689–701.

Chandereng, Thevaa, and Anthony Gitter. 2020. “Lag Penalized Weighted Correlation for Time Series Clustering.” BMC Bioinformatics 21 (1): 21.

Chapman, Tracey, Jenny Bangham, Giovanna Vinti, Beth Seifried, Oliver Lung, Mariana F. Wolfner, Hazel K. Smith, and Linda Partridge. 2003. “The Sex Peptide of *Drosophila Melanogaster*: Female Post-Mating Responses Analyzed by Using RNA Interference.” Proceedings of the National Academy of Sciences of the United States of America 100 (17): 9923–28.

Chatterjee, Nirmalya, and Norbert Perrimon. 2021. “What Fuels the Fly: Energy Metabolism in *Drosophila* and Its Application to the Study of Obesity and Diabetes.” Science Advances. https://doi.org/10.1126/sciadv.abg4336.

Chng, Wen-Bin Alfred, Maroun S. Bou Sleiman, Fanny Schüpfer, and Bruno Lemaitre. 2014. “Transforming Growth Factor β/activin Signaling Functions as a Sugar-Sensing Feedback Loop to Regulate Digestive Enzyme Expression.” Cell Reports 9 (1): 336–48.

Chung, Henry, David W. Loehlin, Héloïse D. Dufour, Kathy Vaccarro, Jocelyn G. Millar, and Sean B. Carroll. 2014. “A Single Gene Affects Both Ecological Divergence and Mate Choice in *Drosophila*.” Science 343 (6175): 1148–51.

Ciofani, Maria, Aviv Madar, Carolina Galan, Maclean Sellars, Kieran Mace, Florencia Pauli, Ashish Agarwal, et al. 2012. “A Validated Regulatory Network for Th17 Cell Specification.” Cell 151 (2): 289–303.

Cognigni, Paola, Andrew P. Bailey, and Irene Miguel-Aliaga. 2011. “Enteric Neurons and Systemic Signals Couple Nutritional and Reproductive Status with Intestinal Homeostasis.” Cell Metabolism 13 (1): 92–104.

Conine, Colin, Fengyun Sun, Lina Song, Jaime A. Rivera-Pérez, Oliver J. Rando. 2018. “Small RNAs Gained during Epididymal Transit of Sperm Are Essential for Embryonic Development in Mice.” Developmental cell, 46(4), 470–480.e3. https://doi.org/10.1016/j.devcel.2018.06.024

Corrigan, Laura, Siamak Redhai, Aaron Leiblich, Shih-Jung Fan, Sumeth M.W. Perera, Rachel Patel, Carina Gandy, Mark S. Wainwright, John F. Morris, Freddie Hamdy, Deborah C.I. Goberdhan, Clive Wilson. 2014. “BMP-Regulated Exosomes from *Drosophila* Male Reproductive Glands Reprogram Female Behavior.” The Journal of cell biology, 206(5), 671–688. https://doi.org/10.1083/jcb.201401072

Csardi, Gabor and Nepusz Tamas (2006). “The igraph Software Package for Complex Network Research.” InterJournal, Complex Systems, 1695. https://igraph.org.

Dalton, Justin E., Tanvi S. Kacheria, Simon Rv Knott, Matthew S. Lebo, Allison Nishitani, Laura E. Sanders, Emma J. Stirling, Ari Winbush, and Michelle N. Arbeitman. 2010. “Dynamic, Mating-Induced Gene Expression Changes in Female Head and Brain Tissues of *Drosophila Melanogaster*.” BMC Genomics 11 (October): 541.

Darlington, T. K., K. Wager-Smith, M. F. Ceriani, D. Staknis, N. Gekakis, T. D. Steeves, C. J. Weitz, J. S. Takahashi, and S. A. Kay. 1998. “Closing the Circadian Loop: CLOCK-Induced Transcription of Its Own Inhibitors per and Tim.” Science 280 (5369): 1599–1603.

Diaz, Fernando, Carson W. Allan, Therese Ann Markow, Jeremy M. Bono, and Luciano M. Matzkin. 2021. “Gene Expression and Alternative Splicing Dynamics Are Perturbed in Female Head Transcriptomes Following Heterospecific Copulation.” BMC Genomics 22 (1): 359.

Ditch, Lynn M., Troy Shirangi, Jeffrey L. Pitman, Kristin L. Latham, Kim D. Finley, Philip T. Edeen, Barbara J. Taylor, and Michael McKeown. 2005. “*Drosophila* Retained/dead Ringer Is Necessary for Neuronal Pathfinding, Female Receptivity and Repression of Fruitless Independent Male Courtship Behaviors.” Development 132 (1): 155–64.

Dobin, Alexander, Carrie A. Davis, Felix Schlesinger, Jorg Drenkow, Chris Zaleski, Sonali Jha, Philippe Batut, Mark Chaisson, and Thomas R. Gingeras. 2013. “STAR: Ultrafast Universal RNA-Seq Aligner.” Bioinformatics 29 (1): 15–21.

Domanitskaya, Elena V., Huanfa Liu, Shanjun Chen, and Eric Kubli. 2007. “The Hydroxyproline Motif of Male Sex Peptide Elicits the Innate Immune Response in *Drosophila* Females.” The FEBS Journal 274 (21): 5659–68.

Donlea, Jeffrey M., Diogo Pimentel, and Gero Miesenböck. 2014. “Neuronal Machinery of Sleep Homeostasis in *Drosophila*.” Neuron 81 (6): 1442.

Donlea, Jeffrey M., Diogo Pimentel, Clifford B. Talbot, Anissa Kempf, Jaison J. Omoto, Volker Hartenstein, and Gero Miesenböck. 2018. “Recurrent Circuitry for Balancing Sleep Need and Sleep.” Neuron. https://doi.org/10.1016/j.neuron.2017.12.016.

Dubowy, Christine and Amita Sehgal. 2017. “Circadian Rhythms and Sleep in *Drosophila melanogaster*.” Genetics 205 (4): 1373–1397.

Eddison, Mark, Douglas J. Guarnieri, Ling Cheng, Che-Hsiung Liu, Kevin G. Moffat, Graeme Davis, and Ulrike Heberlein. 2011. “Arouser Reveals a Role for Synapse Number in the Regulation of Ethanol Sensitivity.” Neuron 70 (5): 979–90.

Erion, Renske, Anna N. King, Gang Wu, John B. Hogenesch, and Amita Sehgal. 2016. “Neural Clocks and Neuropeptide F/Y Regulate Circadian Gene Expression in a Peripheral Metabolic Tissue.” eLife. https://doi.org/10.7554/elife.13552.

Farina, Lorenzo, Alberto De Santis, Samanta Salvucci, Giorgio Morelli, Ida Ruberti. 2008. “Embedding mRNA stability in correlation analysis of time-series gene expression data.” PLoS computational biology, 4(8), e1000141

Feng, Kai, Mark T. Palfreyman, Martin Häsemeyer, Aaron Talsma, and Barry J. Dickson. 2014. “Ascending SAG Neurons Control Sexual Receptivity of *Drosophila* Females.” Neuron 83 (1): 135–48.

Findlay, Geoffrey D., Xianhua Yi, Michael J. MacCoss, Willie J. Swanson. 2008. “Proteomics Reveals Novel *Drosophila* Seminal Fluid Proteins Transferred at Mating.” PLoS biology, 6(7), e178. https://doi.org/10.1371/journal.pbio.0060178

Fischer, David S., Fabian J. Theis, and Nir Yosef. 2018. “Impulse Model-Based Differential Expression Analysis of Time Course Sequencing Data.” Nucleic Acids Research 46 (20): e119.

Fowler, Emily K., Thomas Bradley, Simon Moxon, and Tracey Chapman. 2019. “Divergence in Transcriptional and Regulatory Responses to Mating in Male and Female Fruitflies.” Scientific Reports 9 (1): 16100.

Freeman, M. R., A. Dobritsa, P. Gaines, W. A. Segraves, and J. R. Carlson. 1999. “The Dare Gene: Steroid Hormone Production, Olfactory Behavior, and Neural Degeneration in *Drosophila*.” Development 126 (20): 4591–4602.

Fujii, Shinsuke, and Hubert Amrein. 2002. “Genes Expressed in the *Drosophila* Head Reveal a Role for Fat Cells in Sex-Specific Physiology.” The EMBO Journal 21 (20): 5353–63.

Fulgham, Carson V., Austin P. Dreyer, Anita Nasseri, Asia N. Miller, Jacob Love, Madison M. Martin, Daniel A. Jabr, Sumit Saurabh, and Daniel J. Cavanaugh. 2021. “Central and Peripheral Clock Control of Circadian Feeding Rhythms.” Journal of Biological Rhythms 36 (6): 548–66.

Garaulet, Daniel L., Albertomaria Moro, Eric C. Lai. 2021. “A Double-Negative Regulatory Circuit Underlies the Virgin Behavioral State.” Cell Reports 36 (1): 109335. https://doi.org/10.1016/j.celrep.2021.109335

Garbe, David S., Abigail S. Vigderman, Emilia Moscato, Abigail E. Dove, Christopher G. Vecsey, Matthew S. Kayser, and Amita Sehgal. 2016. “Changes in Female *Drosophila* Sleep Following Mating Are Mediated by SPSN-SAG Neurons.” Journal of Biological Rhythms 31 (6): 551–67.

Gioti, A., S. Wigby, B. Wertheim, E. Schuster, P. Martinez, C. J. Pennington, L. Partridge, and T. Chapman. 2012. “Sex Peptide of *Drosophila Melanogaster* Males Is a Global Regulator of Reproductive Processes in Females.” Proceedings. Biological Sciences / The Royal Society 279 (1746): 4423–32.

Grewal, Savraj S., Ling Li, Amir Orian, Robert N. Eisenman, and Bruce A. Edgar. 2005. “Myc-Dependent Regulation of Ribosomal RNA Synthesis during *Drosophila* Development.” Nature Cell Biology 7 (3): 295–302.

Gu, Zuguang, Roland Eils, and Matthias Schlesner. 2016. “Complex Heatmaps Reveal Patterns and Correlations in Multidimensional Genomic Data.” Bioinformatics 32 (18): 2847–49.

Hao, Yuhan, Stephanie Hao, Erica Andersen-Nissen, William M. Mauck 3rd, Shiwei Zheng, Andrew Butler, Maddie J. Lee, et al. 2021. “Integrated Analysis of Multimodal Single-Cell Data.” Cell 184 (13): 3573–87.e29.

Harshman, L. G., A. M. Loeb, and B. A. Johnson. 1999. “Ecdysteroid Titers in Mated and Unmated *Drosophila Melanogaster* Females.” Journal of Insect Physiology 45 (6): 571–77.

Häsemeyer, Martin, Nilay Yapici, Ulrike Heberlein, and Barry J. Dickson. 2009. “Sensory Neurons in the *Drosophila* Genital Tract Regulate Female Reproductive Behavior.” Neuron 61 (4): 511–18.

Heifetz, Yael, Moshe Lindner, Yuval Garini, and Mariana F. Wolfner. 2014. “Mating Regulates Neuromodulator Ensembles at Nerve Termini Innervating the *Drosophila* Reproductive Tract.” Current Biology: CB 24 (7): 731–37.

Herndon, L. A., and M. F. Wolfner. 1995. “A *Drosophila* Seminal Fluid Protein, Acp26Aa, Stimulates Egg Laying in Females for 1 Day after Mating.” Proceedings of the National Academy of Sciences. https://doi.org/10.1073/pnas.92.22.10114.

Hollis, Brian, Mareike Koppik, Kristina U. Wensing, Hanna Ruhmann, Eléonore Genzoni, Berra Erkosar, Tadeusz J. Kawecki, Claudia Fricke, and Laurent Keller. 2019. “Sexual Conflict Drives Male Manipulation of Female Postmating Responses in *Drosophila Melanogaster*.” Proceedings of the National Academy of Sciences. https://doi.org/10.1073/pnas.1821386116.

Hopkins, Ben R., and Jennifer C. Perry. 2022. “The Evolution of Sex Peptide: Sexual Conflict, Cooperation and Evolution.” Biological reviews of the Cambridge Philosophical Society, 10.1111/brv.12849. Advance online publication. https://doi.org/10.1111/brv.12849

Isaac, R. Elwyn, Chenxi Li, Amy E. Leedale, and Alan D. Shirras. 2010. “*Drosophila* Male Sex Peptide Inhibits Siesta Sleep and Promotes Locomotor Activity in the Post-Mated Female.” Proceedings. Biological Sciences / The Royal Society 277 (1678): 65–70.

Jang, Yong-Hoon, Hyo-Seok Chae, and Young-Joon Kim. 2017. “Female-Specific Myoinhibitory Peptide Neurons Regulate Mating Receptivity in *Drosophila Melanogaster*.” Nature Communications 8 (1): 1630.

Jin, Xi, Yao Tian, Zi Chao Zhang, Pengyu Gu, Chang Liu, and Junhai Han. 2021. “A Subset of DN1p Neurons Integrates Thermosensory Inputs to Promote Wakefulness via CNMa Signaling.” Current Biology: CB 31 (10): 2075–87.e6.

Johnson, Dorothy M., Michael B. Wells, Rebecca Fox, Joslynn S. Lee, Rajprasad Loganathan, Daniel Levings, Abigail Bastien, Matthew Slattery, and Deborah J. Andrew. 2020. “CrebA Increases Secretory Capacity through Direct Transcriptional Regulation of the Secretory Machinery, a Subset of Secretory Cargo, and Other Key Regulators.” Traffic 21 (9): 560–77.

Jo, Kyuri, Inyoung Sung, Dohoon Lee, Hyuksoon Jang, and Sun Kim. 2021. “Inferring Transcriptomic Cell States and Transitions Only from Time Series Transcriptome Data.” Scientific Reports 11 (1): 12566.

Kempf, Anissa, Seoho M. Song, Clifford B. Talbot, and Gero Miesenböck. 2019. “A Potassium Channel β-Subunit Couples Mitochondrial Electron Transport to Sleep.” Nature 568 (7751): 230–34.

Kim, Boram, Makoto I. Kanai, Yangkyun Oh, Minsoo Kyung, Eun-Kyoung Kim, In-Hwan Jang, Ji-Hoon Lee, Sang-Gyu Kim, Greg S. B. Suh, and Won-Jae Lee. 2021. “Response of the Microbiome-Gut-Brain Axis in *Drosophila* to Amino Acid Deficit.” Nature 593 (7860): 570–74.

King-Jones, Kirst, Jean-Philippe Charles, Geanette Lam, and Carl S. Thummel. 2005. “The Ecdysone-Induced DHR4 Orphan Nuclear Receptor Coordinates Growth and Maturation in *Drosophila*.” Cell 121 (5): 773–84.

Kubli, E. 2003. “Sex-Peptides: Seminal Peptides of the *Drosophila* Male.” Cellular and Molecular Life Sciences: CMLS 60 (8): 1689–1704.

Kubli, Eric, and Daniel Bopp. 2012. “Sexual Behavior: How Sex Peptide Flips the Postmating Switch of Female Flies.” Current Biology: CB.

Larkin, Aoife, Steven J. Marygold, Giulia Antonazzo, Helen Attrill, Gilberto dos Santos, Phani V. Garapati, Joshua L. Goodman, L. Sian Gramates, Gillian Millburn, Victor B. Strelets, Christopher J. Tabone, Jim Thurmond J. FlyBase Consortium. 2021. “FlyBase: Updates to the *Drosophila melanogaster* Knowledge Base.” Nucleic Acids Res. 49(D1) D899–D907 DOI:10.1093/nar/gkaa1026

Laturney, Meghan, and Jean-Christophe Billeter. 2016. “*Drosophila Melanogaster* Females Restore Their Attractiveness after Mating by Removing Male Anti-Aphrodisiac Pheromones.” Nature Communications 7 (August): 12322.

Law, Charity W., Yunshun Chen, Wei Shi, and Gordon K. Smyth. 2014. “Voom: Precision Weights Unlock Linear Model Analysis Tools for RNA-Seq Read Counts.” Genome Biology 15 (2): R29.

Leader, David P., Sue A. Krause, Aniruddha Pandit, Shireen A. Davies, and Julian A. T. Dow. 2018. “FlyAtlas 2: A New Version of the *Drosophila Melanogaster* Expression Atlas with RNA-Seq, miRNA-Seq and Sex-Specific Data.” Nucleic Acids Research 46 (D1): D809–15.

Lee, Hyunjin, Hyun Woo Choi, Chen Zhang, Zee-Yong Park, and Young-Joon Kim. 2016. “A Pair of Oviduct-Born Pickpocket Neurons Important for Egg-Laying in *Drosophila Melanogaster*.” Molecules and Cells 39 (7): 573–79.

Lee, Kang-Min, Ivana Daubnerová, R. Elwyn Isaac, Chen Zhang, Sekyu Choi, Jongkyeong Chung, and Young-Joon Kim. 2015. “A Neuronal Pathway That Controls Sperm Ejection and Storage in Female *Drosophila*.” Current Biology: CB 25 (6): 790–97.

Leiblich, Aaron, Luke Marsden, Carina Gandy, Laura Corrigan, Rachel Jenkins, Freddie Hamdy, Clive Wilson. 2012. “Bone Morphogenetic Protein- and Mating-Dependent Secretory Cell Growth and Migration in the *Drosophila* Accessory Gland.” Proceedings of the National Academy of Sciences of the United States of America, 109(47), 19292–19297. https://doi.org/10.1073/pnas.1214517109

Li, Hongjie, Jasper Janssens, Maxime De Waegeneer, Sai Saroja Kolluru, Kristofer Davie, Vincent Gardeux, Wouter Saelens, Fabrice David, Maria Brbić, Jure Leskovec, Colleen N. McLaughlin, Qijing Xie, Robert C. Jones, Katja Brueckner, Jiwon Shim, Sudhir Gopal Tattikota, Frank Schnorrer, Katja Rust, Todd G. Nystul, Zita Carvalho-Santos, Carlos Ribeiro, Soumitra Pal, Teresa M. Przytycka, Aaron M. Allen, Stephen F. Goodwin, Cameron W. Berry, Margaret T. Fuller, Helen White-Cooper, Erika L. Matunis, Stephen DiNardo, Anthony Galenza, Lucy Erin O’Brien, Julian A. T. Dow, FCA Consortium, Heinrich Jasper, Brian Oliver, Norbert Perrimon, Bart Deplancke, Stephen R. Quake, Liqun Luo, Stein Aerts. 2022. “Fly Cell Atlas: a Single-Cell Transcriptomic Atlas of the Adult Fruit Fly.” Science 375 (6584), eabk2432. https://doi.org/10.1126/science.abk2432

Li, Wanhe, Zikun Wang, Sheyum Syed, Cheng Lyu, Samantha Lincoln, Jenno O’Neil, Andrew D. Nguyen, Irena Feng, Michael W. Young. 2021. “Chronic Social Isolation Signals Starvation and Reduces Sleep in *Drosophila*.” Nature 597:239–244.

Liu, Chang, Zhiqiang Meng, Timothy D. Wiggin, Junwei Yu, Martha L. Reed, Fang Guo, Yunpeng Zhang, Michael Rosbash, Leslie C. Griffith. 2019. “A Serotonin-Modulated Circuit Controls Sleep Architecture to Regulate Cognitive Function Independent of Total Sleep in *Drosophila*.” Current Biology 29(21):3635–3646.

Liu, Huanfa, and Eric Kubli. 2003. “Sex-Peptide Is the Molecular Basis of the Sperm Effect in *Drosophila Melanogaster*.” Proceedings of the National Academy of Sciences of the United States of America 100 (17): 9929–33.

Love, Michael I., Wolfgang Huber, and Simon Anders. 2014. “Moderated Estimation of Fold Change and Dispersion for RNA-Seq Data with DESeq2.” Genome Biology 15 (12): 550.

Mack, Paul D., Anat Kapelnikov, Yael Heifetz, and Michael Bender. 2006. “Mating-Responsive Genes in Reproductive Tissues of Female *Drosophila Melanogaster*.” Proceedings of the National Academy of Sciences of the United States of America 103 (27): 10358–63.

Ma, Dingbang, Dariusz Przybylski, Katharine C. Abruzzi, Matthias Schlichting, Qunlong Li, Xi Long, and Michael Rosbash. 2021. “A Transcriptomic Taxonomy of Circadian Neurons around the Clock.” eLife 10 (January). https://doi.org/10.7554/eLife.63056.

Marcillac, Fabrice, François Bousquet, Josiane Alabouvette, Fabrice Savarit, and Jean-François Ferveur. 2005. “A Mutation with Major Effects on *Drosophila Melanogaster* Sex Pheromones.” Genetics 171 (4): 1617–28.

Martelli, Carlotta, Ulrike Pech, Simon Kobbenbring, Dennis Pauls, Britta Bahl, Mirjam Vanessa Sommer, Atefeh Pooryasin, et al. 2017. “SIFamide Translates Hunger Signals into Appetitive and Feeding Behavior in *Drosophila*.” Cell Reports 20 (2): 464–78.

McCarthy, Davis J., Yunshun Chen, and Gordon K. Smyth. 2012. “Differential Expression Analysis of Multifactor RNA-Seq Experiments with Respect to Biological Variation.” Nucleic Acids Research 40 (10): 4288–97.

McDonough-Goldstein, Caitlin E., Kirill Borziak, Scott Pitnick, and Steve Dorus. 2021. “*Drosophila* Female Reproductive Tract Gene Expression Reveals Coordinated Mating Responses and Rapidly Evolving Tissue-Specific Genes.” G3 11 (3). https://doi.org/10.1093/g3journal/jkab020.

McGraw, Lisa A., Andrew G. Clark, and Mariana F. Wolfner. 2008. “Post-Mating Gene Expression Profiles of Female *Drosophila Melanogaster* in Response to Time and to Four Male Accessory Gland Proteins.” Genetics 179 (3): 1395–1408.

McGraw, Lisa A., Greg Gibson, Andrew G. Clark, and Mariana F. Wolfner. 2004. “Genes Regulated by Mating, Sperm, or Seminal Proteins in Mated Female *Drosophila Melanogaster*.” Current Biology. https://doi.org/10.1016/j.cub.2004.08.028.

Mihajlovic, Zorana, Dajana Tanasic, Adam Bajgar, Raquel Perez-Gomez, Pavel Steffal, and Alena Krejci. 2019. “Lime Is a New Protein Linking Immunity and Metabolism in Drosophila.” Developmental Biology 452 (2): 83–94.

Minakuchi, Chieka, Xiaofeng Zhou, and Lynn M. Riddiford. 2008. “Krüppel Homolog 1 (Kr-h1) Mediates Juvenile Hormone Action during Metamorphosis of *Drosophila Melanogaster*.” Mechanisms of Development 125 (1-2): 91–105.

Misra, Snigdha, and Mariana F. Wolfner. 2020. “Seminal Sex Peptide Associates with Rival as Well as Own Sperm, Providing SP Function in Polyandrous Females.” eLife 9 (July). https://doi.org/10.7554/eLife.58322.

Moshitzky, P., I. Fleischmann, N. Chaimov, P. Saudan, S. Klauser, E. Kubli, and S. W. Applebaum. 1996. “Sex-Peptide Activates Juvenile Hormone Biosynthesis in the *Drosophila Melanogaster* Corpus Allatum.” Archives of Insect Biochemistry and Physiology 32 (3-4): 363–74.

Musselman, Laura Palanker, Jill L. Fink, Prasanna Venkatesh Ramachandran, Bruce W. Patterson, Adewole L. Okunade, Ezekiel Maier, Michael R. Brent, John Turk, and Thomas J. Baranski. 2013. “Role of Fat Body Lipogenesis in Protection against the Effects of Caloric Overload in *Drosophila*.” The Journal of Biological Chemistry 288 (12): 8028–42.

Panchal, Trupti, Xi Chen, Ekaterina Alchits, Youjin Oh, James Poon, Jane Kouptsova, Frank A. Laski, and Dorothea Godt. 2017. “Specification and Spatial Arrangement of Cells in the Germline Stem Cell Niche of the *Drosophila* Ovary Depend on the Maf Transcription Factor Traffic Jam.” PLoS Genetics 13 (5): e1006790.

Pascual-Garcia, Pau, Brian Debo, Jennifer R. Aleman, Jessica A. Talamas, Yemin Lan, Nha H. Nguyen, Kyoung J. Won, and Maya Capelson. 2017. “Metazoan Nuclear Pores Provide a Scaffold for Poised Genes and Mediate Induced Enhancer-Promoter Contacts.” Molecular Cell 66 (1): 63–76.e6.

Pasquier, Claude, and Alain Robichon. 2022. “Temporal and Sequential Order of Nonoverlapping Gene Networks Unraveled in Mated Female *Drosophila*.” Life Science Alliance 5 (2). https://doi.org/10.26508/lsa.202101119.

Patke, Alina, Michael W. Young, and Sofia Axelrod. 2020. “Molecular Mechanisms and Physiological Importance of Circadian Rhythms.” Nature Reviews. Molecular Cell Biology 21 (2): 67–84.

Peng, Jing, Shanjun Chen, Susann Büsser, Huanfa Liu, Thomas Honegger, and Eric Kubli. 2005. “Gradual Release of Sperm Bound Sex-Peptide Controls Female Postmating Behavior in *Drosophila*.” Current Biology: CB 15 (3): 207–13.

Peng, Jing, Peder Zipperlen, and Eric Kubli. 2005. “*Drosophila* Sex-Peptide Stimulates Female Innate Immune System after Mating via the Toll and Imd Pathways.” Current Biology: CB 15 (18): 1690–94.

Perry, Jennifer C., Laura Sirot, Stuart Wigby. 2013. “The Seminal Symphony: How to Compose an Ejaculate.” Trends in ecology & evolution, 28(7), 414–422. https://doi.org/10.1016/j.tree.2013.03.005

Pimentel, Diogo, Jeffrey M. Donlea, Clifford B. Talbot, Seoho M. Song, Alexander J. F. Thurston, and Gero Miesenböck. 2016. “Operation of a Homeostatic Sleep Switch.” Nature 536 (7616): 333–37.

Pilch, Bartosz & Matthias Mann. 2006. “Large-Scale and High-Confidence Proteomic Analysis of Human Seminal Plasma.” Genome biology, 7(5), R40. https://doi.org/10.1186/gb-2006-7-5-r40

Poiani, Aldo. 2006. “Complexity of Seminal Fluid: a Review.” Behavioral Ecology and Sociobiology 60: 289–310.

Prokupek, A. M., S. D. Kachman, I. Ladunga, and L. G. Harshman. 2009. “Transcriptional Profiling of the Sperm Storage Organs of *Drosophila* Melanogaster.” Insect Molecular Biology 18 (4): 465–75.

Reiff, Tobias, Jake Jacobson, Paola Cognigni, Zeus Antonello, Esther Ballesta, Kah Junn Tan, Joanne Y. Yew, Maria Dominguez, and Irene Miguel-Aliaga. 2015. “Endocrine Remodelling of the Adult Intestine Sustains Reproduction in *Drosophila*.” eLife 4 (July): e06930.

Ren, Guilin R., Frank Hauser, Kim F. Rewitz, Shu Kondo, Alexander F. Engelbrecht, Anders K. Didriksen, Suzanne R. Schjøtt, et al. 2015. “CCHamide-2 Is an Orexigenic Brain-Gut Peptide in *Drosophila*.” PloS One 10 (7): e0133017.

Rezával, Carolina, Tetsuya Nojima, Megan C. Neville, Andrew C. Lin, and Stephen F. Goodwin. 2014. “Sexually Dimorphic Octopaminergic Neurons Modulate Female Postmating Behaviors in *Drosophila*.” Current Biology: CB 24 (7): 725–30.

Rezával, Carolina, Hania J. Pavlou, Anthony J. Dornan, Yick-Bun Chan, Edward A. Kravitz, and Stephen F. Goodwin. 2012. “Neural Circuitry Underlying *Drosophila* Female Postmating Behavioral Responses.” Current Biology: CB 22 (13): 1155–65.

Ribeiro, Carlos, and Barry J. Dickson. 2010. “Sex Peptide Receptor and Neuronal TOR/S6K Signaling Modulate Nutrient Balancing in *Drosophila*.” Current Biology: CB 20 (11): 1000–1005.

Robinson, Mark D., Davis J. McCarthy, and Gordon K. Smyth. 2010. “edgeR: A Bioconductor Package for Differential Expression Analysis of Digital Gene Expression Data.” Bioinformatics 26 (1): 139–40.

Rubinstein, C. Dustin, and Mariana F. Wolfner. 2013. “*Drosophila* Seminal Protein Ovulin Mediates Ovulation through Female Octopamine Neuronal Signaling.” Proceedings of the National Academy of Sciences of the United States of America 110 (43): 17420–25.

Sakai, T., and N. Ishida. 2001. “Circadian Rhythms of Female Mating Activity Governed by Clock Genes in *Drosophila*.” Proceedings of the National Academy of Sciences of the United States of America 98 (16): 9221–25.

Saudan, Philippe, Klaus Hauck, Matthias Soller, Yves Choffat, Michael Ottiger, Michael Spörri, Zhaobing Ding, et al. 2002. “Ductus Ejaculatorius Peptide 99B (DUP99B), a Novel *Drosophila Melanogaster* Sex-Peptide Pheromone.” European Journal of Biochemistry / FEBS 269 (3): 989–97.

Sayols, Sergi. 2020. “rrvgo: a Bioconductor Package to Reduce and Visualize Gene Ontology Terms.” https://ssayols.github.io/rrvgo.

Scheffer, Louis K., Shan C. Xu, Michal Januszewski., Zhiyuan Lu, Shin-Ya Takemura, Kenneth J. Hayworth, Gary B. Huang, Kazunori Shinomiya, Jeremy Maitlin-Shepard, Stuart Berg, Jody Clements, Philip M. Hubbard, William T. Katz, Lowell Umayam, Ting Zhao, David Ackerman, Tim Blakely, John Bogovic, Tom Dolafi, Dagmar Kainmueller, … Stephen M. Plaza. 2020. “A Connectome and Analysis of the Adult *Drosophila* Central Brain.” eLife, 9, e57443. https://doi.org/10.7554/eLife.57443.

Scheunemann, L., A. Lampin-Saint-Amaux, J. Schor, and T. Preat. 2019. “A Sperm Peptide Enhances Long-Term Memory in Female *Drosophila*.” Science Advances. https://doi.org/10.1126/sciadv.aax3432.

Schlamp, Florencia, Sofie Y. N. Delbare, Angela M. Early, Martin T. Wells, Sumanta Basu, and Andrew G. Clark. 2021. “Dense Time-Course Gene Expression Profiling of the *Drosophila Melanogaster* Innate Immune Response.” BMC Genomics 22 (1): 304.

Scott, D. 1986. “Sexual Mimicry Regulates the Attractiveness of Mated *Drosophila Melanogaster* Females.” Proceedings of the National Academy of Sciences of the United States of America 83 (21): 8429–33.

Shao, Lisha, Phuong Chung, Allan Wong, Igor Siwanowicz, Clement F. Kent, Xi Long, and Ulrike Heberlein. 2019. “A Neural Circuit Encoding the Experience of Copulation in Female *Drosophila*.” Neuron 102 (5): 1025–36.e6.

Sokolowski, Dustin J., Mariela Faykoo-Martinez, Lauren Erdman, Huayun Hou, Cadia Chan, Helen Zhu, Melissa M. Holmes, Anna Goldenberg, and Michael D. Wilson. 2021. “Single-Cell Mapper (scMappR): Using scRNA-Seq to Infer the Cell-Type Specificities of Differentially Expressed Genes.” NAR Genomics and Bioinformatics. https://doi.org/10.1093/nargab/lqab011.

Soller, Matthias, Irmgard U. Haussmann, Martin Hollmann, Yves Choffat, Kalpana White, Eric Kubli, and Mireille A. Schäfer. 2006. “Sex-Peptide-Regulated Female Sexual Behavior Requires a Subset of Ascending Ventral Nerve Cord Neurons.” Current Biology: CB 16 (18): 1771–82.

Soneson, Charlotte, Katarina L. Matthes, Malgorzata Nowicka, Charity W. Law, and Mark D. Robinson. 2016. “Isoform Prefiltering Improves Performance of Count-Based Methods for Analysis of Differential Transcript Usage.” Genome Biology 17 (January): 12.

Sun, Jinghan, Chang Liu, Xiaobing Bai, Xiaoting Li, Jingyun Li, Zhiping Zhang, Yunpeng Zhang, Jing Guo, and Yan Li. 2017. “*Drosophila* FIT Is a Protein-Specific Satiety Hormone Essential for Feeding Control.” Nature Communications 8 (January): 14161.

Szklarczyk, Damian, Annika L. Gable, David Lyon, Alexander Junge, Stefan Wyder, Jaime Huerta-Cepas, Milan Simonovic, et al. 2019. “STRING v11: Protein-Protein Association Networks with Increased Coverage, Supporting Functional Discovery in Genome-Wide Experimental Datasets.” Nucleic Acids Research 47 (D1): D607–13.

Taylor, Sean C., Katia Nadeau, Meysam Abbasi, Claude Lachance, Marie Nguyen, and Joshua Fenrich. 2019. “The Ultimate qPCR Experiment: Producing Publication Quality, Reproducible Data the First Time.” Trends in Biotechnology 37 (7): 761–74.

Terhzaz, Selim, Philippe Rosay, Stephen F. Goodwin, and Jan A. Veenstra. 2007. “The Neuropeptide SIFamide Modulates Sexual Behavior in *Drosophila*.” Biochemical and Biophysical Research Communications 352 (2): 305–10.

The Gene Ontology Consortium. 2021. “The Gene Ontology Resource: Enriching a GOld Mine.” Nucleic Acids Research 49(D1):D325–D334

Thompson, Jeffrey K., Matthew R. Peterson, and Ralph D. Freeman. 2003. “Single-Neuron Activity and Tissue Oxygenation in the Cerebral Cortex.” Science 299 (5609): 1070–72.

Tram, Uyen., and Mariana. F. Wolfner. 1998. “Seminal Fluid Regulation of Female Sexual Attractiveness in *Drosophila Melanogaster*.” Proceedings of the National Academy of Sciences of the United States of America 95 (7): 4051–54.

Uchizono, Shun, Yumi Tabuki, Natsumi Kawaguchi, Teiichi Tanimura, and Taichi Q. Itoh. 2017. “Mated *Drosophila Melanogaster* Females Consume More Amino Acids during the Dark Phase.” PloS One 12 (2): e0172886.

Venkatraman, Sara, Sumanta Basu, Andrew G. Clark, Sofie Delbare, Myung Hee Lee, and Martin T. Wells. n.d. “An Empirical Bayes Approach to Estimating Dynamic Models of Co-Regulated Gene Expression.” https://doi.org/10.1101/2021.07.08.451684.

Wainwright, Mark S., Ben R. Hopkins, Cláudia C. Mendes, Aashika Sekar, Benjamin Kroeger, Josephine E. E. U. Hellberg, Shih-Jung Fan, Abigail Pavey, Pauline P. Marie, Aaron Leiblich, Irem Sepil, Philip D. Charles, Marie L. Thézénas, Roman Fischer, Benedikt M. Kessler, Carina Gandy, Laura Corrigan, Rachel Patel, Stuart Wigby, John F. Morris, Deborah C. I. Goberdhan, Clive Wilson. 2021. “*Drosophila* Sex Peptide Controls the Assembly of Lipid Microcarriers in Seminal Fluid. Proceedings of the National Academy of Sciences of the United States of America, 118(5) e2019622118. https://doi.org/10.1073/pnas.2019622118

Walker, Samuel James, Verónica María Corrales-Carvajal, Carlos Ribeiro. 2015. “Postmating Circuitry Modulates Salt Taste Processing to Increase Reproductive Output in *Drosophila*.” Current Biology: CB 25 (20): 2621–30.

Wang, Fei, Kaiyu Wang, Nora Forknall, Christopher Patrick, Tansy Yang, Ruchi Parekh, Davi Bock, and Barry J. Dickson. 2020. “Neural Circuitry Linking Mating and Egg Laying in *Drosophila* Females.” Nature 579 (7797): 101–5.

Wang, Kaiyu, Fei Wang, Nora Forknall, Tansy Yang, Christopher Patrick, Ruchi Parekh, and Barry J. Dickson. 2021. “Neural Circuit Mechanisms of Sexual Receptivity in *Drosophila* Females.” Nature 589 (7843): 577–81.

White, Melissa A., Alessandro Bonfini, Mariana F. Wolfner, and Nicolas Buchon. 2021. “*Drosophila Melanogaster* Sex Peptide Regulates Mated Female Midgut Morphology and Physiology.” Proceedings of the National Academy of Sciences. https://doi.org/10.1073/pnas.2018112118.

Wicker-Thomas, C., C. Henriet, and R. Dallerac. 1997. “Partial Characterization of a Fatty Acid Desaturase Gene in *Drosophila Melanogaster*.” Insect Biochemistry and Molecular Biology 27 (11): 963–72.

Wickham, Hadley. 2009. “ggplot2: Elegant Graphics for Data Analysis.” Springer Science & Business Media.

Wigby, Stuart, Tracey Chapman. 2005. “Sex Peptide Causes Mating Costs in Female Drosophila melanogaster.” Current biology: CB, 15(4), 316–321. https://doi.org/10.1016/j.cub.2005.01.051

Wilkinson, Leland. 2012. “Exact and Approximate Area-Proportional Circular Venn and Euler Diagrams.” IEEE Transactions on Visualization and Computer Graphics 18 (2): 321–31.

Wu, Gang, Ron C. Anafi, Michael E. Hughes, Karl Kornacker, and John B. Hogenesch. 2016. “MetaCycle: An Integrated R Package to Evaluate Periodicity in Large Scale Data.” Bioinformatics 32 (21): 3351–53.

Wu, Tianzhi, Erqiang Hu, Shuangbin Xu, Meijun Chen, Pingfan Guo, Zehan Dai, Tingze Feng, et al. 2021. “clusterProfiler 4.0: A Universal Enrichment Tool for Interpreting Omics Data.” Innovation (New York, N.Y.) 2 (3): 100141.

Xiao, Shuke, Jennifer S. Sun, and John R. Carlson. 2019. “Robust Olfactory Responses in the Absence of Odorant Binding Proteins.” eLife 8 (October). https://doi.org/10.7554/eLife.51040.

Xu, Kanyan, Justin R. DiAngelo, Michael E. Hughes, John B. Hogenesch, and Amita Sehgal. 2011. “The Circadian Clock Interacts with Metabolic Physiology to Influence Reproductive Fitness.” Cell Metabolism 13 (6): 639–54.

Xu, Kanyan, Xiangzhong Zheng, and Amita Sehgal. 2008. “Regulation of Feeding and Metabolism by Neuronal and Peripheral Clocks in *Drosophila*.” Cell Metabolism 8 (4): 289–300.

Yang, Chung-Hui, Sebastian Rumpf, Yang Xiang, Michael D. Gordon, Wei Song, Lily Y. Jan, and Yuh-Nung Jan. 2009. “Control of the Postmating Behavioral Switch in *Drosophila* Females by Internal Sensory Neurons.” Neuron 61 (4): 519–26.

Yapici, Nilay, Young-Joon Kim, Carlos Ribeiro, and Barry J. Dickson. 2008. “A Receptor That Mediates the Post-Mating Switch in *Drosophila* Reproductive Behavior.” Nature 451 (7174): 33–37.

Yu, Guangchuang, Li-Gen Wang, Yanyan Han, and Qing-Yu He. 2012. “clusterProfiler: An R Package for Comparing Biological Themes Among Gene Clusters.” OMICS: A Journal of Integrative Biology. https://doi.org/10.1089/omi.2011.0118.

Zhang, Chen, Ivana Daubnerova, Yong-Hoon Jang, Shu Kondo, Dušan Žitňan, and Young-Joon Kim. 2021. “The Neuropeptide Allatostatin C from Clock-Associated DN1p Neurons Generates the Circadian Rhythm for Oogenesis.” Proceedings of the National Academy of Sciences. https://doi.org/10.1073/pnas.2016878118.

Zhang, Yuqing, Giovanni Parmigiani, and W. Evan Johnson. 2020. “: Batch Effect Adjustment for RNA-Seq Count Data.” NAR Genomics and Bioinformatics 2 (3): lqaa078.

Zhou, Chuan, Yufeng Pan, Carmen C. Robinett, Geoffrey W. Meissner, and Bruce S. Baker. 2014. “Central Brain Neurons Expressing Doublesex Regulate Female Receptivity in *Drosophila*.” Neuron 83 (1): 149–63.

Zhu, Anqi, Joseph G. Ibrahim, and Michael I. Love. 2019. “Heavy-Tailed Prior Distributions for Sequence Count Data: Removing the Noise and Preserving Large Differences.” Bioinformatics 35 (12): 2084–92.

Zipper L, Denise Jassmann, Sofie Burgmer, Bastian Görlich, and Tobias Reiff. 2020. “Ecdysone Steroid Hormone Remote Controls Intestinal Stem Cell Fate Decisions via the PPARγ- Homolog Eip75B in *Drosophila*.” eLife. https://doi.org/10.7554/elife.55795.

